# Focused Ultrasound Blood-Brain Barrier Opening Arrests the Growth and Formation of Cerebral Cavernous Malformations

**DOI:** 10.1101/2024.01.31.577810

**Authors:** Delaney G. Fisher, Khadijeh A. Sharifi, Ishaan M. Shah, Catherine M. Gorick, Victoria R. Breza, Anna C. Debski, Matthew R. Hoch, Tanya Cruz, Joshua D. Samuels, Jason P. Sheehan, David Schlesinger, David Moore, John R. Lukens, G. Wilson Miller, Petr Tvrdik, Richard J. Price

## Abstract

**BACKGROUND:** Cerebral cavernous malformations (CCM) are vascular lesions within the central nervous system, consisting of dilated and hemorrhage-prone capillaries. CCMs can cause debilitating neurological symptoms, and surgical excision or stereotactic radiosurgery are the only current treatment options. Meanwhile, transient blood-brain barrier opening (BBBO) with focused ultrasound (FUS) and microbubbles is now understood to exert potentially beneficial bioeffects, such as stimulation of neurogenesis and clearance of amyloid-β. Here, we tested whether FUS BBBO could be deployed therapeutically to control CCM formation and progression in a clinically-representative murine model.

**METHODS:** CCMs were induced in mice by postnatal, endothelial-specific *Krit1* ablation. FUS was applied for BBBO with fixed peak-negative pressures (PNPs; 0.2-0.6 MPa) or passive cavitation detection-modulated PNPs. Magnetic resonance imaging (MRI) was used to target FUS treatments, evaluate safety, and measure longitudinal changes in CCM growth after BBBO.

**RESULTS:** FUS BBBO elicited gadolinium accumulation primarily at the perilesional boundaries of CCMs, rather than lesion cores. Passive cavitation detection and gadolinium contrast enhancement were comparable in CCM and wild-type mice, indicating that *Krit1* ablation does not confer differential sensitivity to FUS BBBO. Acutely, CCMs exposed to FUS BBBO remained structurally stable, with no signs of hemorrhage. Longitudinal MRI revealed that FUS BBBO halted the growth of 94% of CCMs treated in the study. At 1 month, FUS BBBO-treated lesions lost, on average, 9% of their pre-sonication volume. In contrast, non-sonicated control lesions grew to 670% of their initial volume. Lesion control with FUS BBBO was accompanied by a marked reduction in the area and mesenchymal appearance of *Krit* mutant endothelium. Strikingly, in mice receiving multiple BBBO treatments with fixed PNPs, *de novo* CCM formation was significantly reduced by 81%. Mock treatment plans on MRIs of patients with surgically inaccessible lesions revealed their lesions are amenable to FUS BBBO with current clinical technology.

**CONCLUSIONS:** Our results establish FUS BBBO as a novel, non-invasive modality that can safely arrest murine CCM growth and prevent their *de novo* formation. As an incisionless, MR image-guided therapy with the ability to target eloquent brain locations, FUS BBBO offers an unparalleled potential to revolutionize the therapeutic experience and enhance the accessibility of treatments for CCM patients.

## Introduction

Cerebral cavernous malformations (CCM) are vascular lesions originating in the capillary-venous vessels of the central nervous system^1^. These slow flow vascular malformations are hemorrhage prone, grossly enlarged, and lack many of the supporting cells of the neurovascular unit^2,3^. CCMs generally arise due to biallelic mutation in one of the three CCM-related genes: *Krit1/CCM1*, *MGC4607/CCM2*, and *PDCD10/CCM3*^1,4^. CCM patients can experience debilitating and life-altering symptoms such as motor and visual deficits, seizures, and stroke^5^. These symptoms generally arise from the rapid growth and hemorrhage of a CCM^6^. The current standard of care for CCM is invasive surgical resection. However, resection is associated with a high risk of post-operative morbidities and limited to surgically accessible CCMs^6^. Due to their eloquent location, CCMs in the brainstem are associated with even greater risks of early morbidity and recurrent growth following incomplete resection^6,7^. Stereotactic radiosurgery is also a treatment option but conveys risks associated with ionizing radiation that can lead to adverse radiation effects^8^. The pathological trajectory of CCMs remains largely uncertain to clinicians^9–11^. Thus, CCM patients, and parents of children with CCM, are put in the position of choosing between the risks of neurosurgery or inaction.

As an incisionless therapy with the ability to target eloquent brain locations, focused ultrasound (FUS) may represent an ideal alternative for CCM treatment. With targeting provided by magnetic resonance imaging (MRI), FUS delivers acoustic energy deep within the body to non-invasively produce mechanical or thermal therapeutic effects^12^. When FUS is combined with an intravenous (i.v.) injection of gas-filled microbubbles, the oscillating pressure waves induce an alternating expansion and contraction of the gas within microbubbles, which in turn causes the microbubbles to push and pull on the walls of blood vessels. If performed in the brain, this procedure can induce a temporary opening of the blood-brain barrier (BBB).

FUS-mediated BBB opening (BBBO) has been deployed primarily to enable enhanced delivery of drugs and other therapeutic agents into the brain for various neurological conditions^13–15^. However, FUS BBBO has also been shown to be beneficial in the absence of drug delivery for the treatment of Alzheimer’s disease^16–22^. While the exact mechanism(s) behind the beneficial effect of FUS BBBO in Alzheimer’s disease are not completely understood, ample preclinical evidence of this effect has led to several clinical trials that are testing this approach in patients with Alzheimer’s disease (NCT04118764, NCT04526262, NCT02986932, NCT03739905, NCT04250376). In this study, we examined the effectiveness and safety profile of FUS BBBO applied to CCMs and its potential to, in the absence of drug delivery, therapeutically control the growth and *de novo* formation of CCMs.

## Results

### FUS effectively opens the BBB within the CCM microenvironment

Given the altered biomechanical properties^23–25^ and increased caliber of the vasculature of CCMs and the surrounding perilesional vasculature (**Figure 1A**), we first questioned whether FUS in combination with i.v. microbubble injection could effectively elicit BBBO in CCM mice. We acquired baseline, high resolution T2-weighted spin echo MR images of CCM mice to select CCMs for sonication. On the day of FUS treatment, gadolinium contrast agent (gadobenate dimeglumine; 1.058 kDa) was injected intravenously, and a pre-sonication T1-weighted spin echo MR image was obtained. We next performed FUS BBBO on selected CCMs using peak-negative pressures (PNP), i.e. ultrasound wave amplitudes, of 0.2 MPa - 0.6 MPa and standard BBBO parameters. Analysis of the T1 contrast enhancement revealed that FUS BBBO enhanced gadolinium accumulation to the CCM (**Figure 1B-C)**. Gadolinium accumulation around CCMs was significantly increased by FUS BBBO over the baseline leakiness of gadolinium for PNPs of 0.3 MPa to 0.6 MPa (**Figure 1C**) and primarily localized to the perilesional boundaries of the sonicated CCM, rather than the lesion core (**Figure 1B)**. Thus, FUS can effectively open the BBB within the CCM microenvironment, despite the enlarged and irregular microvasculature associated with the lesion.

**Figure 1.**
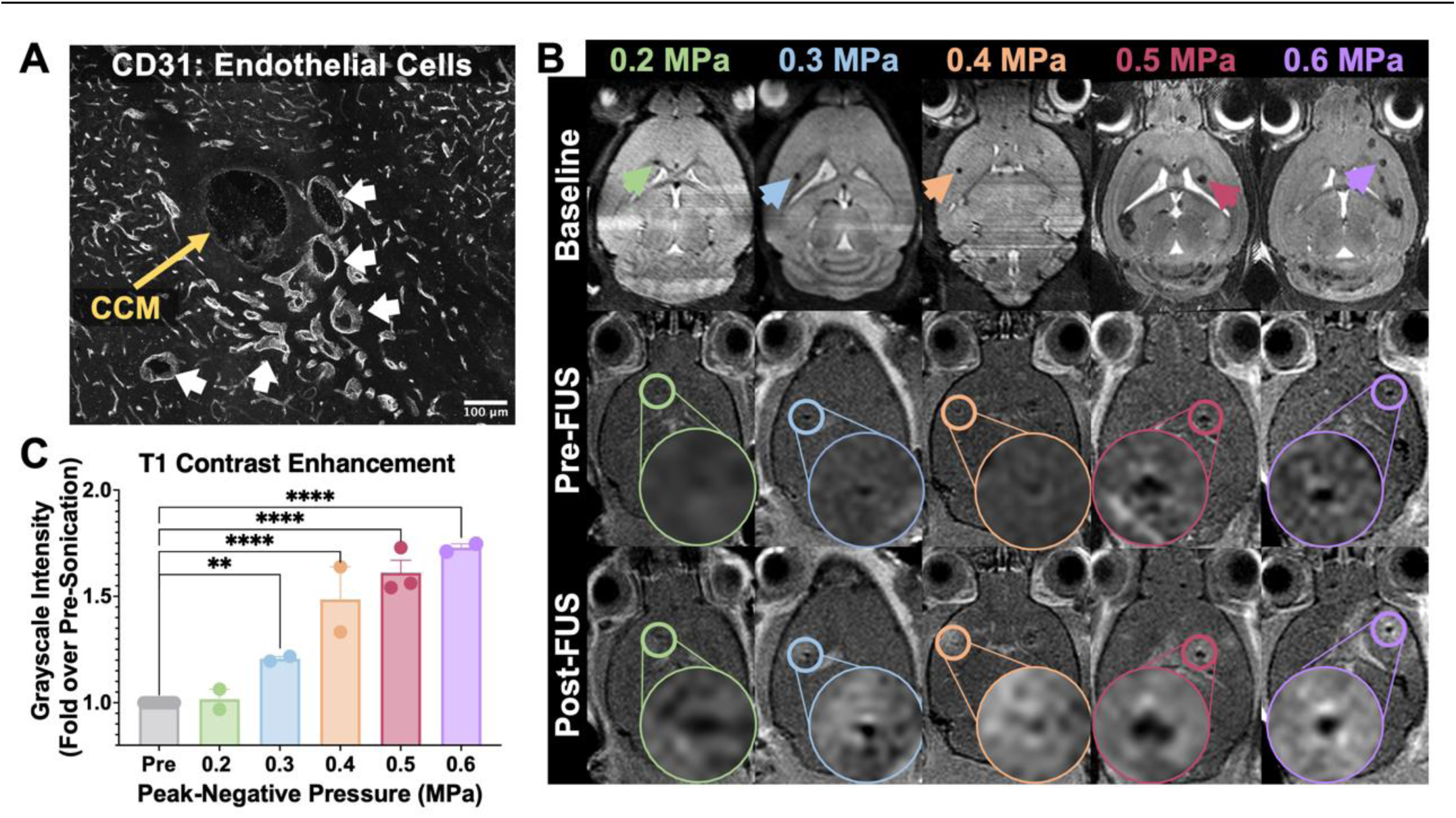
FUS effectively opens the BBB within the CCM microenvironment. (A) Confocal image of a CCM (in the absence of FUS) stained with CD31 for endothelial cells. Image depicts the grossly enlarged CCM core (yellow arrow), and moderately dilated perilesional vasculature (white arrows). Scale bar = 100 µm. (B) Top row: Baseline, high-resolution T2-weighted spin echo images used for selecting CCMs for FUS targeting. Arrowheads indicate selected CCMs. Middle row: T1-weighted spin echo images acquired following gadolinium contrast agent injection but immediately prior to FUS application. Circles indicate targeted CCMs, and insets display magnified views of the targeted CCMs. Bottom row: T1-weighted spin echo images acquired following gadolinium contrast agent injection and FUS application. Columns indicate PNPs used for sonication. T1 contrast enhancement is visible following FUS BBBO and localized to perilesional boundaries of the sonicated CCM. (C) Bar graph of T1 contrast enhancement quantified as the fold change in grayscale intensity of sonicated CCMs in the post-image over the pre-image (as seen in A). Gadolinium accumulation following FUS BBBO over the baseline CCM leakiness for PNPs of 0.3 MPa to 0.6 MPa. p=0.0054 for 0.3 MPa and p<0.0001 for 0.4 MPa – 0.6 MPa, one-way ANOVA followed by Dunnett’s multiple comparisons test.

### FUS BBBO does not increase volume or bleeding of hemorrhage-prone CCMs acutely

Due to the propensity of CCMs to hemorrhage and, more broadly, the dysregulated state of the microvasculature in CCMs^1^, we next sought to evaluate the safety of FUS BBBO in this disease model. To determine if growth or bleeding was acutely induced by FUS BBBO at PNPs of 0.2 MPa – 0.6 MPa, MR images of the brains of CCM mice were taken before and 24 h after FUS BBBO. A 3-dimensional, T2-weighted spin echo sequence was employed to accurately capture changes in CCM volume (**Figure 2A**), while 3-dimensional, susceptibility-weighted images (SWI) were acquired to capture changes in iron content and fluid flow (i.e. bleeding or hemorrhage; **Figure 2C**) with high sensitivity. Measurement of the hypointense lesion margins between pre- and post-sonication images revealed no evidence of acute growth or hemorrhage induced by FUS BBBO (**Figure 2B, D**), indicating that FUS BBBO causes neither growth nor bleeding of CCMs at acute time points. Immunofluorescent staining of erythrocytes with Ter119 (**Figure S1**) confirmed that FUS BBBO did not exacerbate lesion hemorrhage.

**Figure 2.**
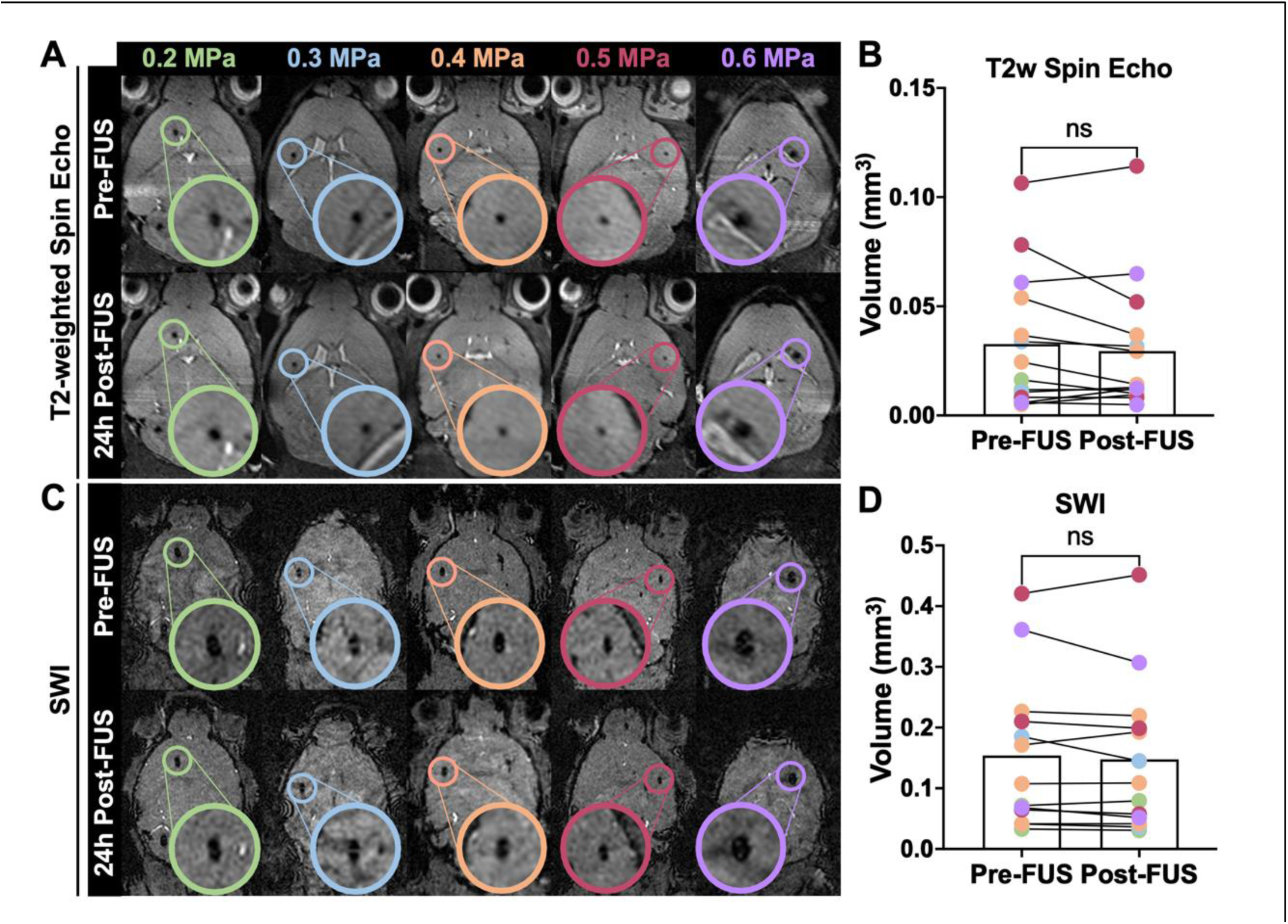
Acute stability of CCMs exposed to FUS BBBO. (A) High-resolution T2-weighted spin echo images displaying either CCMs prior to sonication (top row) or 24 h following sonication (bottom row). Circles denote targeted CCMs, and insets display magnified views of the targeted CCMs. (B) Targeted CCM volumes prior to sonication and 24 h following sonication on T2-weighted spin echo images with color indicating applied PNP. CCM volume does not significantly demonstrate changes in volume following sonication. p=0.41, Wilcoxon matched-pairs signed rank test. (C) High-resolution susceptibility-weighted images of the same mice in A, displaying either CCMs prior to sonication (top row) or 24 h following sonication (bottom row). (D) Targeted CCM volumes prior to sonication and 24 h following sonication on susceptibility-weighted images with color indicating applied PNP. CCM volume does not significantly demonstrate changes in bleeding following sonication. p=0.34, Wilcoxon matched-pairs signed rank test.

### Comparison of FUS BBBO contrast enhancement and acoustic emission signatures between wild-type and CCM mice

To test whether CCM mice differentially respond to FUS BBBO at PNPs of 0.4 MPa – 0.6 MPa, we compared T1 contrast enhancement, which is indicative of the degree of BBBO and contrast delivery, and passive cavitation detection (PCD) measurements, which is indicative of the microbubble activity during sonication, between wild-type mice and CCM mice. Our analysis revealed no significant differences in T1 contrast enhancement between wild-type and CCM mice at any of the tested PNPs (**Figure 3A-B**), suggesting that the extent of BBBO is comparable. To compare the microbubble activity, spectrograms of the frequency response for each burst during the FUS application were generated (**Figure 3C**), and cavitation levels were quantified for spectra signifying unstable and stable microbubble activity (**Figure 3D-E**). Spectral domains associated with a transition towards or an increase in unstable, inertial cavitation of microbubbles (i.e. subharmonic, ultraharmonics, and broadband)^26,27^ increased with PNP and were comparable between wild-type and CCM mice (**Figure 3D**). Spectral domains associated with stable cavitation (i.e. harmonics)^27,28^ were comparable for PNPs of 0.4 MPa and 0.5 MPa (**Figure 3E**). However, at a PNP of 0.6 MPa, CCM mice displayed an increase in harmonic emissions, while the harmonic emissions of wild-type mice remained similar to that observed at lower PNPs (**Figure 3E**). Altogether, these results suggest that FUS BBBO affects wild-type and CCM mice similarly with regards to the degree of BBBO and microbubble activity induced, particularly unstable microbubble activity. Meanwhile, at high PNPs, stable microbubble activity is enhanced in CCM mice, albeit without comparable increases in unstable, inertial cavitation.

**Figure 3.**
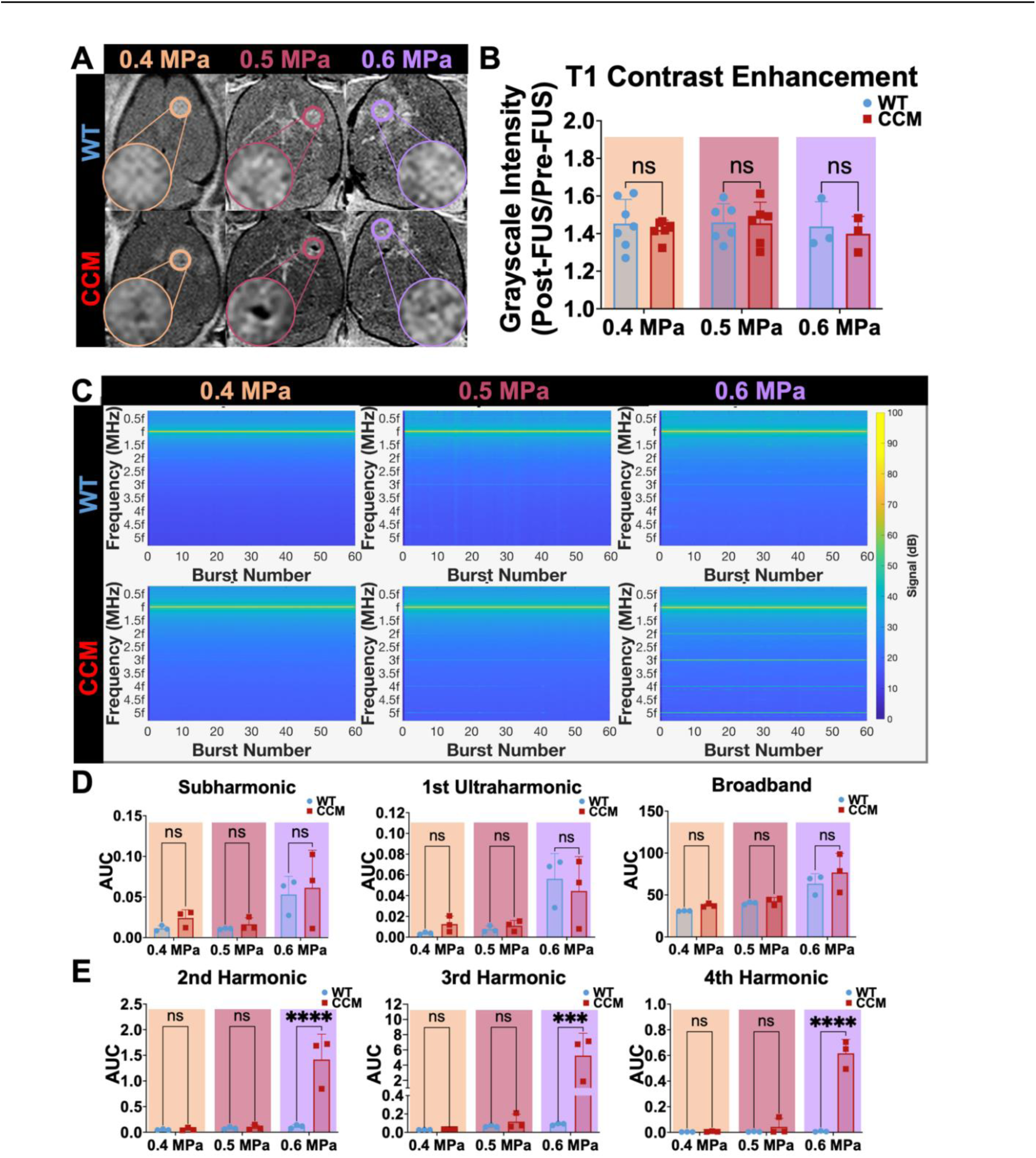
Comparison of FUS BBBO contrast enhancement and acoustic emission signatures between wild-type and CCM mice. (A) Representative T1-weighted spin echo images acquired following gadolinium contrast agent injection and FUS application in wild-type mice or CCM mice for PNPs of 0.4 MPa – 0.6 MPa. (B) Bar graph of T1 contrast enhancement. Enhancement is comparable in wild-type and CCM mice for PNPs of 0.4 MPa-0.6 MPa. p=0.92 for 0.4 MPa, p=0.9998 for 0.5 MPa, and p=0.96 for 0.6 MPa; two-way ANOVA with Šidák’s multiple comparison test. (C) Spectrograms of the frequency response for each burst during the FUS application averaged over cohorts of wild-type and CCM mice at PNPs of 0.4 MPa – 0.6 MPa (n=3 mice per group and 2-3 sonication replicates per mouse). (D) Subharmonic, first ultraharmonic, and broadband emissions for wild-type and CCM mice at PNPs of 0.4 MPa – 0.6 MPa. p>0.4 for all PNPs, two-way ANOVA with Šidák’s multiple comparisons test. (E) Second, third, and fourth harmonic emissions for wild-type and CCM mice at PNPs of 0.4 MPa – 0.6 MPa, indicating that stable cavitation-associated signatures between wild-type and CCM mice are comparable at 0.4 MPa and 0.5 MPa, but not significantly increased in CCM mice at 0.6 MPa. P > 0.7 for 0.4 – 0.5 MPa and 2nd – 4th harmonics; p < 0.0001, p = 0.0006, p<0.0001 for 0.6 MPa and 2nd, 3rd, and 4th harmonics, respectively; two-way ANOVA with Šidák’s multiple comparisons test.

### CCM mice are not differentially sensitive to adverse effects generated by FUS BBBO at high PNPs

To assess the longitudinal safety of FUS BBBO in CCM mice, we collected T2-weighted spin echo sequences over a one-month period following FUS BBBO in wild-type and CCM mice (**Figure 4A**). Different FUS BBBO regimens were tested: a single FUS BBBO application or repeat applications performed three times for PNPs of 0.4 MPa or two times for PNPs of 0.5 MPa and 0.6 MPa, with a three-day spacing between sonications. Edema, visible as hyperintensity on T2-weighted MRI, was apparent in lesion-free brain tissue in a fraction of both wild-type and CCM mice one day post-FUS BBBO for PNPs of 0.5 MPa and 0.6 MPa (**Figure 4A-B**). Hemosiderin deposits, visible as hypointensity on T2-weighted MRI, were also apparent in lesion-free brain tissue in wild-type and CCM mice at time points beyond one day post-FUS BBBO and persisted for at least one month following FUS BBBO for PNPs of 0.5 MPa and 0.6 MPa (**Figure 4A, C**). Edema, quantified by an increase in the ipsilateral-to-contralateral grayscale ratio, primarily occurred after BBBO with PNPs of 0.5 MPa (**Figure 4B**), and hemosiderin deposition, quantified by a decrease in the ipsilateral-to-contralateral grayscale ratio, increased as a function on PNP (**Figure 4C**). Generally, acute edema was associated with chronic hemosiderin deposition for both models and both treatment arms (**Figure 4D**). When comparing the prevalence of edema and hemosiderin deposition between wild-type and CCM mice for each treatment regimen and PNP, no significant differences were seen (**Figure 4E**). However, when treatment regimens were aggregated, wild-type mice actually exhibited a greater propensity for edema than CCM mice (**Figure 4B**), yet wild-type and CCM mice shared an equivalent correlation for hemosiderin deposition (**Figure 4C**). These results suggest that, while BBBO with PNPs greater than 0.4 MPa are safe for CCMs, FUS BBBO at increased PNPs can induce edema and hemosiderin deposition, consistent with that seen in wild-type mice.

**Figure 4.**
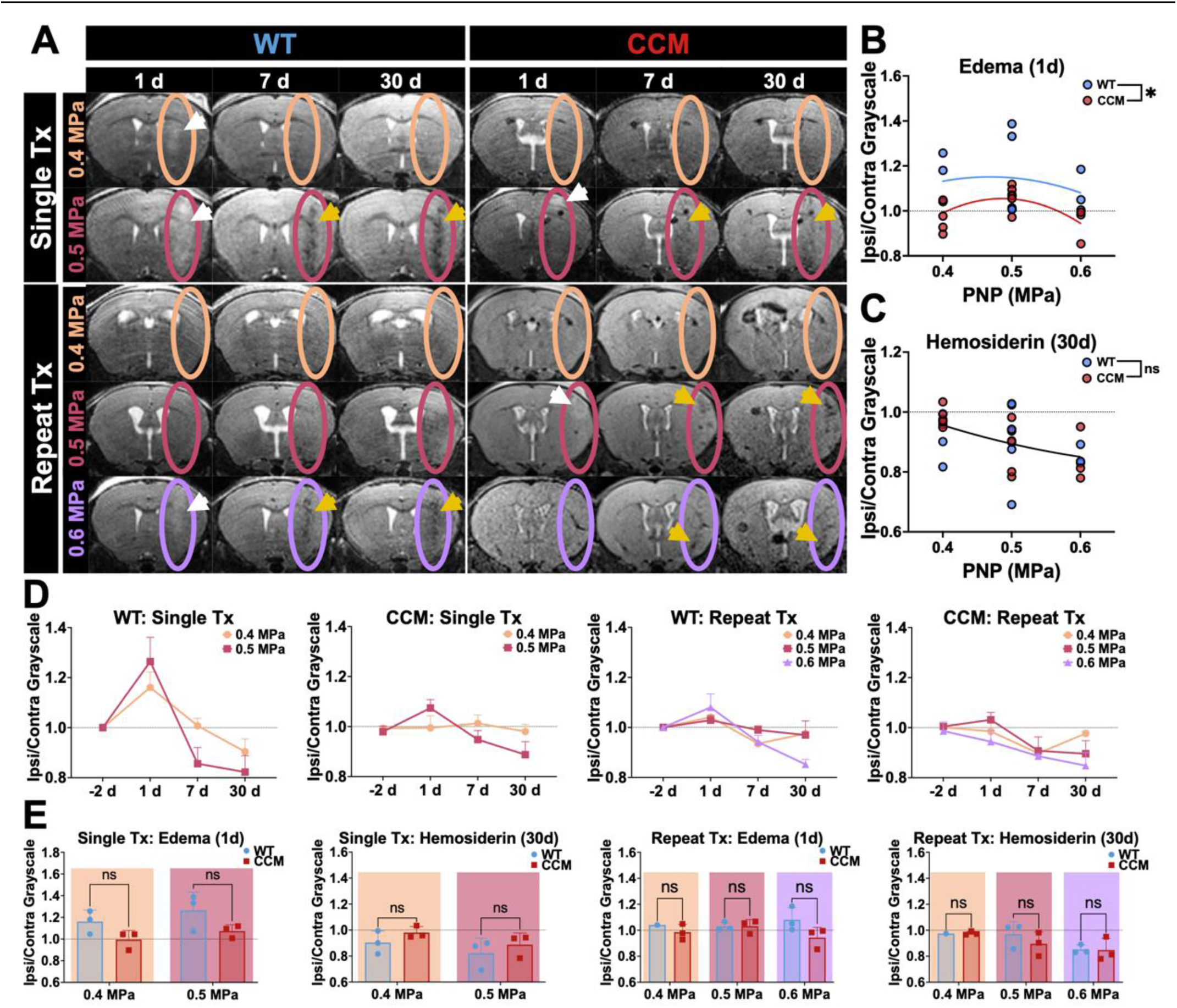
CCM mice are not differentially sensitive to adverse effects generated by FUS BBBO at high PNPs. (A) Representative high resolution, T2-weighted spin echo images of wild-type and CCM mice at 1 d, 7 d, and 30 d post-sonication at PNPs of 0.4 MPa – 0.6 MPa in either a single sonication or repeat sonication treatment regimen. Ovals denote focal column. White arrows denote hyperintensities associated with edema. Yellow arrows denote hypointensities associated with hemosiderin deposition. (B) Scatterplot of ipsilateral-to-contralateral grayscale intensity at 1d post-FUS (when edema is visible) of wild-type and CCM mice for PNPs of 0.4 MPa – 0.6 MPa. p=0.047, comparison of fits with F-test for a 2^nd^ order polynomial regression. (C) Scatterplot of ipsilateral-to-contralateral grayscale intensity at 30d post-FUS (when hemosiderin is visible) of wild-type and CCM mice for PNPs of 0.4 MPa – 0.6 MPa. p=0.77, comparison of fits with F-test for a 2^nd^ order polynomial regression. (D) Line graphs of ipsilateral-to-contralateral grayscale intensities over the one-month imaging period for all PNPs within a mouse model and treatment arm, revealing that edema on day 1 is generally followed by hemosiderin on days 7 and 30. (E) Ipsilateral-to-contralateral grayscale intensities over the one-month imaging period for all PNPs within a mouse model and treatment arm, indicating no significant differences when comparing models at individual PNPs within a treatment arm. p = 0.1368 and p = 0.5386 for both PNPs in the single treatment arm for edema and hemosiderin, respectively; p > 0.7 for PNPs of 0.4 MPa and 0.5 MPa and p = 0.0923 for PNP of 0.6 MPa in the repeat treatment arm for edema; p > 0.5 for all PNPs in the repeat treatment arm for hemosiderin; two-way ANOVA with Holm-Šidák’s multiple comparisons test.

### Real-time PCD-modulation of PNP ensures the safety of sonicated brain tissue without compromising gadolinium delivery

To ensure safety of our FUS BBBO application and examine the effect of more clinically-representative FUS BBBO regimens in CCM mice, we performed FUS BBBO using a real-time PCD feedback control system to modulate the applied PNP during sonication^29–31^. Using this PCD-modulated PNP approach, the maximum PNP occurred within the first 15 seconds of treatment, and the PNP generally decreased gradually over the sonication period (**Figure 5A**). This approach resulted in a time-averaged PNP ranging from 0.23 MPa – 0.30 MPa and a maximum PNP ranging from 0.25 MPa – 0.38 MP. PCD-modulated PNPs successfully increased T1 contrast enhancement in the CCM microenvironment (**Figure 5B-C**). Comparing PCD-modulation of PNP to the fixed PNP approach revealed that PCD-modulated PNP resulted in higher T1 contrast enhancement than fixed PNPs of similar amplitudes (**Figure 5D**). Acoustic emissions measurements revealed that PCD-modulated PNP elicits comparable subharmonic, broadband, and harmonic spectra when compared to fixed PNPs of 0.4 MPa and 0.5 MPa (**Figure 5E-F**). Longitudinal T2-weighted MRI also demonstrated that PCD-modulated PNP obviates edema and hemosiderin deposition following FUS BBBO (**Figure 5G-H**). For BBBO in CCM mice, edema was comparable across PNPs and a reduction of hemosiderin deposition was seen with PNPs averaging less than or equal to 0.4 MPa (**Figure 5I**). Altogether, these data indicate that PCD-modulation of PNP ensures the safety of FUS BBBO in CCM brain tissue and elicits enhanced gadolinium delivery compared to fixed PNPs.

**Figure 5.**
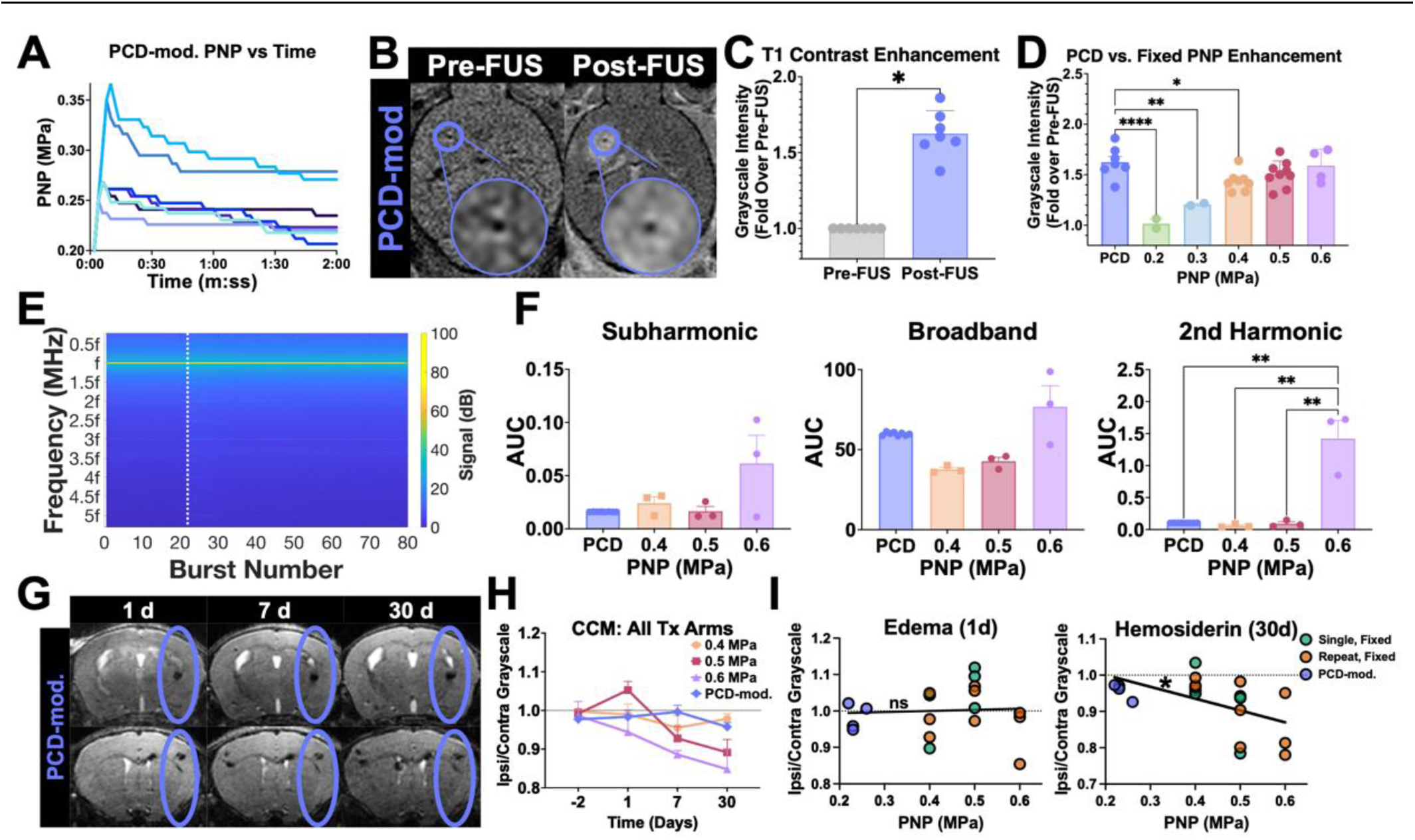
Real-time PCD-modulation of PNP ensures the safety of sonicated brain tissue without compromising gadolinium delivery. (A) Applied PNP versus time during PCD feedback-controlled approach. Each line indicates the average applied PNP across two sonication targets for the same mouse during a single FUS sonication period. (B) Representative T1-weighted contrast images before and after FUS BBBO with PCD-modulated PNPs. (C) Bar graph of T1 contrast enhancement quantified as the fold change in grayscale intensity of sonicated CCMs in the post-image over the pre-image (as seen in B), indicating successful BBBO. p=0.016, Wilcoxon matched-pairs signed rank test. (D) Bar graph of T1 contrast enhancement quantified as the fold change in grayscale intensity of sonicated CCMs in the post-image over the pre-image for CCM mice with fixed PNP and PCD-modulated PNP cohorts. Graphs reveal that T1 contrast enhancement is greater with PCD-modulated PNP compared to fixed PNP in the same range of applied PNP of 0.2 – 0.4 MPa. p < 0.0001 for PCD vs. 0.2 MPa, p = 0.0018 for PCD vs. 0.3 MPa, p = 0.0368 for PCD vs. 0.4 MPa, p = 0.2864 for PCD vs. 0.5 MPa, and p = 0.9918 for PCD vs. 0.6 MPa, one-way ANOVA with Dunnett’s multiple comparison’s test. (E) Spectrogram of the frequency response for each burst during the FUS application averaged over CCM mice with PCD-modulated PNP (n=4 mice and 2 sonication replicates per mouse). Dotted line denotes separation of baseline sonications without microbubbles and sonications with microbubbles. (F) Subharmonic, broadband, and second harmonic emissions for CCM mice at PCD-modulated PNP and fixed PNPs of 0.4 MPa – 0.6 MPa, indicating comparable acoustic signatures for PNPs less than 0.6 MPa. p > 0.8 for the subharmonic, ultraharmonic, and 2^nd^-3^rd^ harmonic emissions for PCD vs. 0.4 or 0.5 MPa; p > 0.3 for the broadband emissions; p = 0.003 for 2^nd^ harmonic emissions and 0.6 MPa vs. PCD,0.4 MPa, and 0.5 MPa; two-way ANOVA with Šidák’s multiple comparisons test. (G) Representative high resolution, T2-weighted spin echo images of wild-type and CCM mice at 1 d, 7 d, and 30 d post-sonication at PNPs of 0.4 MPa – 0.6 MPa in either a single sonication or repeat sonication treatment regimen. Ovals denote focal column. (H) Line graphs of ipsilateral-to-contralateral grayscale intensities over the one-month imaging period for CCM mice and all PNP regimens. (I) Scatterplot of ipsilateral-to-contralateral grayscale intensity versus time-averaged PNP for CCM with single treatments and fixed PNP, repeat treatments and fixed PNP, or repeat treatments and PCD-modulated PNP mice on day 1 (left) or day 30 post-FUS (right). For edema, ipsilateral-to-contralateral grayscale intensity is <u>not</u> significantly correlated with PNP; however, for hemosiderin, ipsilateral-to-contralateral grayscale intensity is significantly correlated with PNP. p = 0.8382 for edema and p = 0.0163 for hemosiderin, linear regression with F test.

### FUS BBBO arrests CCM growth

We then asked if FUS BBBO stimulates therapeutically beneficial responses for CCMs. First, we tested several FUS BBBO regimens for their ability to control the growth of CCMs. CCM mice were placed in (i) a single FUS BBBO regimen with fixed PNP (i.e. one FUS BBBO treatment at either 0.4 MPa or 0.5 MPa), (ii) a repeat FUS BBBO regimen with fixed PNP (i.e. three FUS BBBO treatments at 0.4 MPa or two FUS BBBO treatments at 0.5 MPa or 0.6 MPa, all staged three days apart), or (iii) a repeat FUS BBBO regimen with PCD-modulated PNP (i.e. two FUS BBBO treatments staged three days apart). Mice were treated between 2 and 3 months of age, a period of rapidly escalating lesion burden^32^. Male and female mice across 9 litters were used (**Table S1**), and MR images were acquired following each sonication and up to one month thereafter (**Figure 6A, C, E**). Sonicated CCM volumes were compared to non-sonicated CCMs of similar baseline size and anatomical location within the same cohort of mice. The average sonicated and non-sonicated CCM volume prior to FUS application was 0.039 mm^3^ for both conditions. Remarkably, CCMs exposed to FUS BBBO in all treatment regimens exhibited nearly complete cessation of growth (**Figure 6B, D, F**). Only 3 of 47 CCMs exposed to FUS BBBO grew more than 0.02 mm^3^ in 1 month, while 26 of 41 CCMs not exposed to FUS BBBO grew this amount in the same period. Significant differences in lesion volume between the sonicated and non-sonicated CCMs were seen after 30 days for all treatment arms (**Figure 6B, D, F).** At 7 days, sonicated CCMs were significantly smaller than non-sonicated CCMs in the repeat FUS and fixed PNP arm (**Figure 6D**). At 30 days post-FUS BBBO, sonicated CCMs in all treatment arms demonstrated a markedly reduced mean lesion volume, reaching just 28%, 10%, and 26% of the mean volume of the non-sonicated CCM volume in the single, fixed PNP; repeat, fixed PNP; and repeat, PCD-modulated PNP arms, respectively. Increases in PNP and number of FUS BBBO treatments were both inversely correlated with increased lesion volume (**Figure S2A-B**). The effect of sex on CCM volume and FUS BBBO was also evaluated (**Figure S3A-B**). After 30 days, CCMs in male mice were larger than those in female mice, regardless of FUS BBBO treatment (**Figure S3A, Table S2**). However, sex did not affect the ability of FUS BBBO to control CCM growth (**Figure S3A, Table S2**).

**Figure 6.**
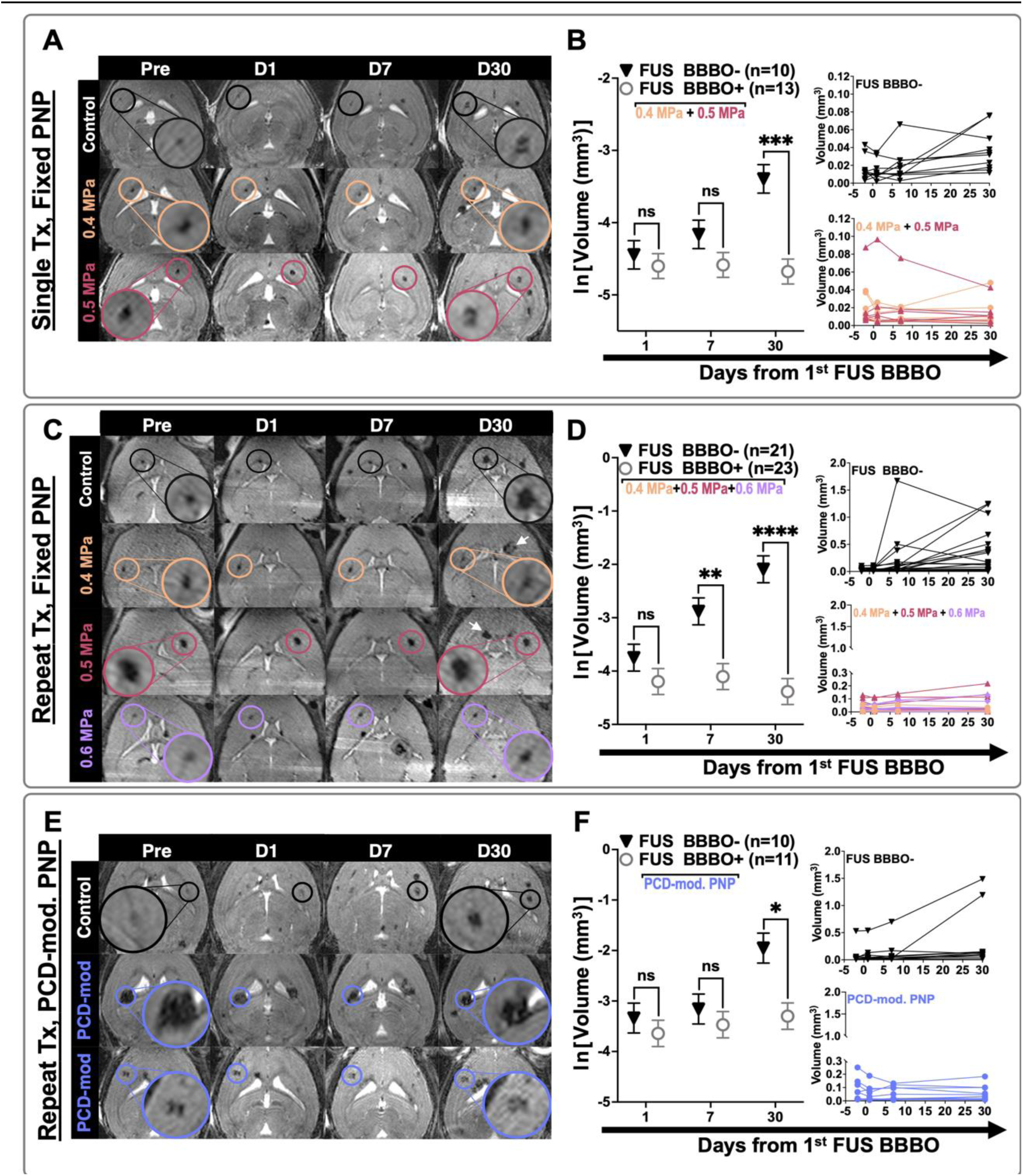
FUS BBBO arrests the growth of CCMs. (A, C, E) Longitudinal T2-weighted spin echo images for representative mice in the (A) single sonication with fixed PNP arm, (C) repeat sonication with fixed PNP arm, or (E) repeat sonication with PCD-modulated PNP arm. Black circles indicate non-sonicated, control lesions, and colored circles indicate sonicated lesions corresponding to PNP applied. White arrows denote new lesions formed in non-sonicated hemisphere. (B, D, F) Left: Summary plots comparing the natural log transform of CCM volume between sonicated CCMs and non-sonicated CCMs for mice in the (B) single sonication with fixed PNP arm, (D) repeat sonication with fixed PNP arm, or (F) repeat sonication with PCD-modulated PNP arm. Right: Line graphs of CCM volume for individual CCMs for each treatment group. At 30 days, sonicated CCMs are significantly smaller than non-sonicated control CCMs for all treatment arms. p = 0.0002, p < 0.0001, and p = 0.0131 for the single, fixed PNP; repeat, fixed PNP; and repeat, PCD-mod. PNP arms, respectively; linear mixed effect model and pairwise comparison with Tukey’s adjustment. At 7 days, sonicated CCMs are significantly smaller than non-sonicated CCMs in the repeat FUS and fixed PNP arm. p = 0.0021, linear mixed effect model and pairwise comparison with Tukey’s adjustment.

### FUS BBBO with fixed PNP and repeat sonications can prevent *de novo* lesion formation

To then ascertain if FUS BBBO impacts the formation of new lesions, we counted the number of lesions contained within the focal zone (i.e. T1-contrast-enhanced brain region) in MR images taken prior to FUS BBBO, as well as one month following FUS BBBO. The same analysis was performed in the contralateral hemisphere of each mouse using the same volume and mirrored anatomical location (**Figure 7A, C, E**). The change in the number of lesions from the pre-image to the 30-days post-FUS BBBO image was compared for the sonicated and contralateral brain areas within each mouse. This analysis revealed that the repeat FUS regimen with fixed PNP significantly reduced the formation of new CCMs by 81% compared to the contralateral brain region (**Figure 7D**). Meanwhile, the single FUS with fixed PNP regimen and repeat FUS with PCD-modulated PNP regimen displayed trends toward reduced *de novo* CCM formation (**Figure 7B, F**). Importantly, in all treatment arms, FUS BBBO did *not* induce an increase in lesion formation. In fact, both the single and repeat FUS with fixed PNP cohorts contained one mouse that displayed fewer lesions in the sonicated brain region one month following FUS BBBO compared to the pre-image, suggesting that some CCMs may be cleared with FUS BBBO. Increases in PNP were found to be significantly, inversely correlated with *de novo* lesion formation, while the number of sonication treatments followed this trend, albeit not significantly (**Figure S2C-D**). The effect of sex on *de novo* CCMs and FUS BBBO was also evaluated (**Figure S3C-D**). Sex did not significantly alter the ability of FUS BBBO to control CCM formation (**Figure S3C, Table S2**).

**Figure 7.**
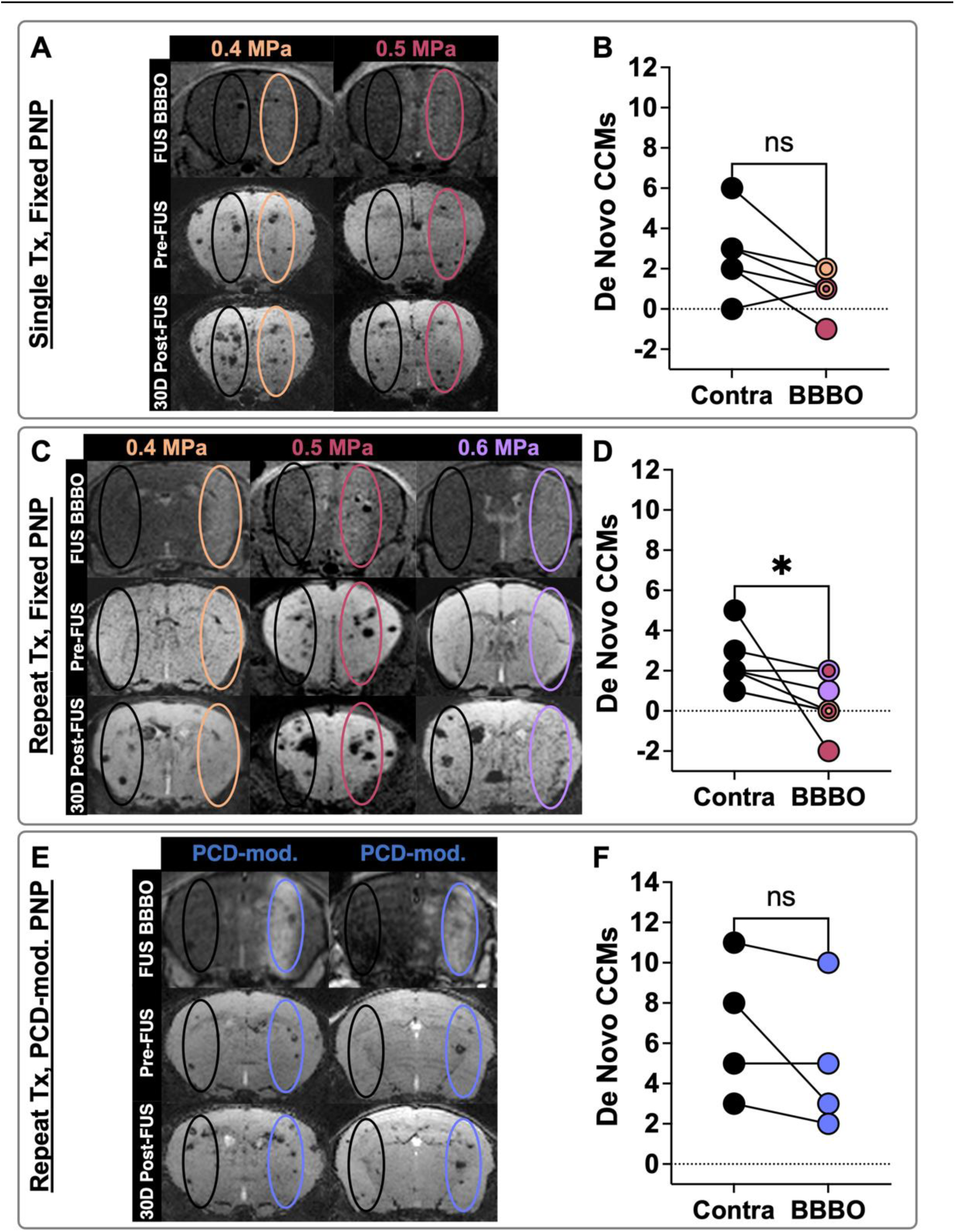
FUS BBBO with fixed PNP and repeat sonications can prevent *de novo* lesion formation. (A, C, E) Top row: T1-weighted spin echo images taken immediately following FUS BBBO with hyperintense signal denoting the focal column. Middle and bottom rows: minimum intensity projection images of longitudinal T2-weighted spin echo images to visualize through 1 mm of the focal column for representative mice in the (A) single sonication with fixed PNP arm, (C) repeat sonication with fixed PNP arm, or (E) repeat sonication with PCD-modulated PNP arm. Black ovals denote contralateral, non-sonicated ROIs for *de novo* quantification, while colored ovals represent sonicated ROIs. (B, D, F) Paired line graphs comparing the change in CCM number one month following FUS BBBO between the sonicated brain region and the contralateral non-sonicated brain region for mice in the (B) single sonication with fixed PNP arm, (D) repeat sonication with fixed PNP arm, or (F) repeat sonication with PCD-modulated PNP arm. Concentric circles indicate multiple mice with the same number of *de novo* CCMs. Colors indicate applied PNP. For mice receiving the repeat FUS regimen with fixed PNP, the number of new lesions formed in the sonicated brain region is significantly reduced compared to the contralateral brain region. p = 0.0312, Wilcoxon matched-pairs signed rank test.

### FUS BBBO restores endothelial morphology to the mutated CCM vasculature and remodels CCM immune landscape

To elucidate how FUS BBBO may halt CCM growth and prevent new lesion formation, we performed an extensive immunohistological analysis of brain sections at 1 day, 7 days, or 30 days post-FUS BBBO. We first questioned whether FUS BBBO affects the *Krit1* mutant endothelium. After the induction of endothelial *Krit1* knock out (*Krit1*KO) in our CCM mouse model, tdTomato is expressed, allowing visualization of the mutated CCM vasculature (**Figure 8A**). As expected, in non-sonicated lesions, the *Krit1*KO vasculature was mesenchymal in appearance and aggressively growing (**Figure 8B**). *Krit1*KO mutant vessel size was comparable between sonicated and non-sonicated lesions at 1 day and 7 days post-FUS BBBO. However, at 30 days post-FUS BBBO, the mesenchymal appearance of *Krit1*KO vasculature underwent a striking reversal to a more endothelial-like morphology (**Figure 8A**). Further, the average area of *Krit1*KO vasculature was significantly reduced (**Figure 8B**), despite no change in the proliferation of *Krit1*KO cells (**Figure S4A-B**).

**Figure 8.**
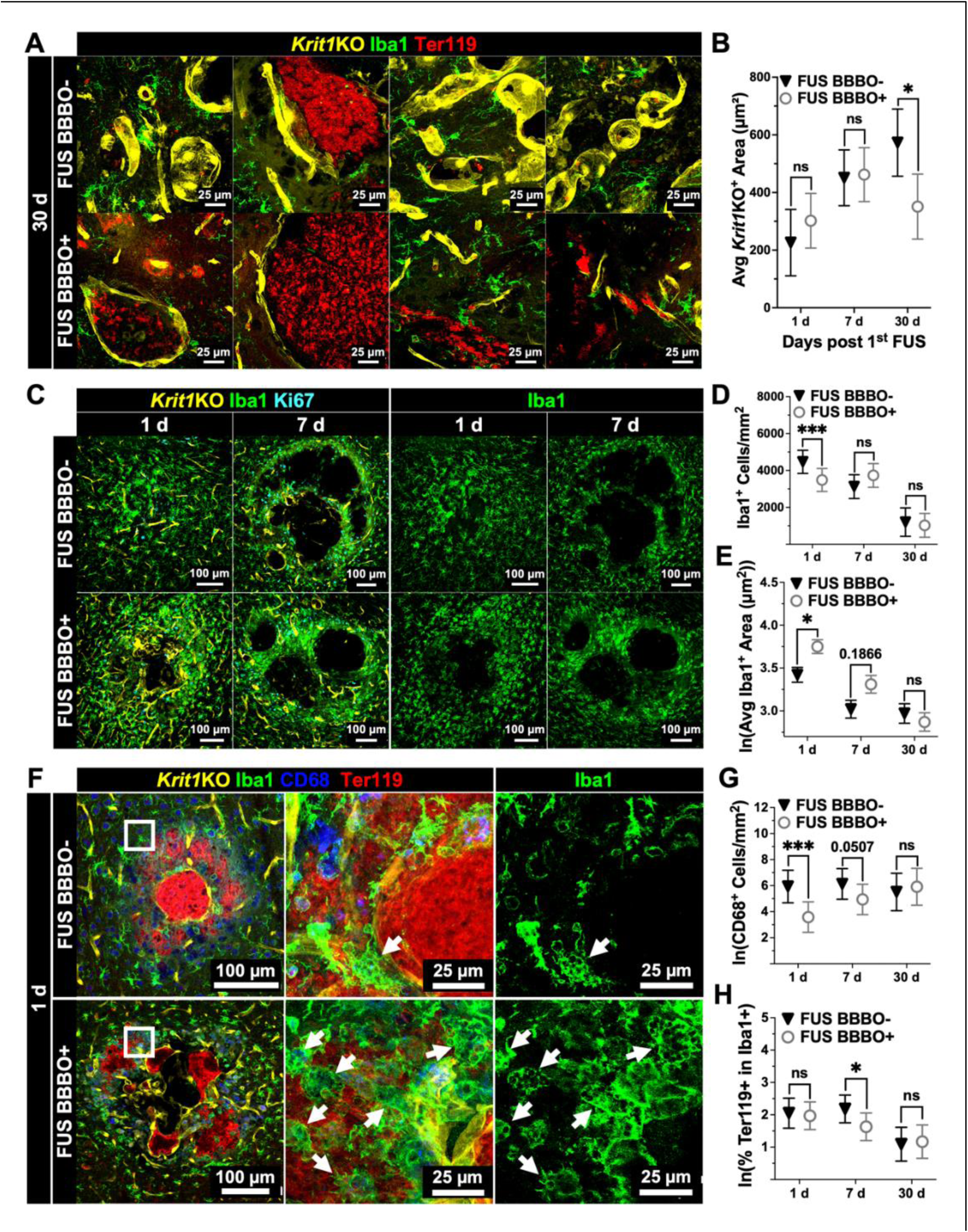
FUS BBBO restores endothelial morphology to the mutated CCM vasculature and remodels CCM immune landscape. (A) Immunofluorescent images of non-sonicated and sonicated CCMs at 30 d post-FUS BBBO with staining for mutated vasculature *(Krit1*KO), microglia/macrophages (Iba1), and erythrocytes (Ter119). The mutated vasculature in sonicated CCMs had reduced mesenchymal appearance compared to non-sonicated CCMs. (B) Graph of average mutated CCM vasculature area at 1 d, 7 d, and 30 d post-FUS BBBO for non-sonicated and sonicated CCMs, indicating reduced area in sonicated CCMs at 30 d. p = 0.0199, linear mixed effect model and pairwise comparison with Tukey’s adjustment. (C) Immunofluorescent images of non-sonicated and sonicated CCMs at 1 d and 7 d post-FUS BBBO with staining for mutated vasculature *(Krit1*KO), microglia/macrophages (Iba1), and proliferation (Ki67). (D) Graph of density of microglia/macrophages at 1 d, 7 d, and 30 d post-FUS BBBO for non-sonicated and sonicated CCMs, revealing a reduced number in sonicated lesions at 1 d. p = 0.0003, linear mixed effects model and pairwise comparison with Tukey’s adjustment. (E) Graph of the natural log of the average microglia/macrophage area at 1 d, 7 d, and 30 d post-FUS BBBO for non-sonicated and sonicated CCMs, demonstrating an increase in microglia/macrophage size in sonicated lesions at 1 d. p = 0.0106, linear mixed effect model and pairwise comparison with Tukey’s adjustment. (F) Immunofluorescent images of non-sonicated and sonicated CCMs at 1 d post-FUS BBBO with staining for mutated vasculature *(Krit1*KO), microglia/macrophages (Iba1), lysosomes (CD68), and erythrocytes (Ter119). Insets display 63x maximum intensity projections of the corresponding 20x image. Arrows denote foamy macrophages. (G) Graph of the natural log of phagocyte density at 1 d, 7 d, and 30 d post-FUS BBBO for non-sonicated and sonicated CCMs, revealing a reduced number in sonicated lesions at 1 d. p = 0.0009, linear mixed effects model and pairwise comparison with Tukey’s adjustment. (H) Graph of the natural log of the percent of erythrocytes colocalized in microglia/macrophages at 1 d, 7 d, and 30 d post-FUS BBBO for non-sonicated and sonicated CCMs, indicating a smaller amount in sonicated lesions at 7 d. p = 0.0303, linear mixed effects model and pairwise comparison with Tukey’s adjustment.

Because FUS BBBO is thought to augment microglial phagocytosis^33,34^, we also looked for evidence of enhanced microglia/macrophage phagocytic activity in sonicated lesions, with particular emphasis on the potential for clearance of erythrocytes. At 1 day post-FUS, the number of Iba1+ cells (microglia/macrophages) was significantly decreased in sonicated lesions (**Figure 8C-D**); however, their average area was significantly increased (**Figure 8C, E-F**). Closer examination revealed these enlarged Iba1+ cells as foamy macrophages (**Figure 8F**). Unexpectedly, the number of cells expressing the phagolysosomal marker CD68 was actually decreased at 1 day and 7 days in sonicated lesions (**Figure 8G**). Further, the percent of red blood cells (Ter119+) colocalized with Iba1+ cells, which would be suggestive of phagocytosis of erythrocytes, was not increased by FUS BBBO. In fact, this metric was actually decreased at 7 days after FUS BBBO (**Figure 8H**). Interestingly, the CD68+ cell population steadily recovers after the acute reduction by FUS BBBO (**Figure 8G**). The proliferation of Iba1+ cells and the proliferation, number, and size of GFAP+ astrocytes were not significantly different between sonicated and non-sonicated lesions at any time point following FUS BBBO (**Figure S4A, C-F**). Finally, we found that CD45+ immune cell infiltration was significantly elevated 7 days post-FUS BBBO in sonicated lesions (**Figure S5A-B**). Inspecting the morphology and location of CD45+, Iba1+, and *Krit1*KO signal revealed monocytes in the lumens of lesions, Iba1+ microglia/macrophage processes extending to CD45+ immune cells, and CD45+Iba1+ cells lining mutated vessels (**Figure S5A**).

### Current clinical FUS systems are equipped to treat CCMs in patients

Finally, to assess the feasibility of clinical CCM treatments with FUS BBBO, we designed FUS BBBO treatment plans for 3 CCM patients with surgically inaccessible CCMs who had instead received stereotactic radiosurgery (SRS)^8^ (**Figure 9**). SRS treatment plans are shown in **Figure 9A**, with 12.5 Gy and 6.3 Gy isodose lines circumscribing the target CCM and its margin. We reimagined these treatment plans for FUS BBBO using the NaviFUS clinical MRI-guided FUS system (**Figure 9B**). These CCMs in eloquent brain locations were accessible for FUS BBBO treatment. A total of 43 sonication points spanning 2 cm in diameter and 8.65 cm^3^ in volume provided adequate coverage of the target CCM in all 3 patients. Thus, we demonstrate that current clinical FUS systems are equipped to treat CCMs in patients, especially those that are not candidates for traditional surgical excision.

**Figure 9.**
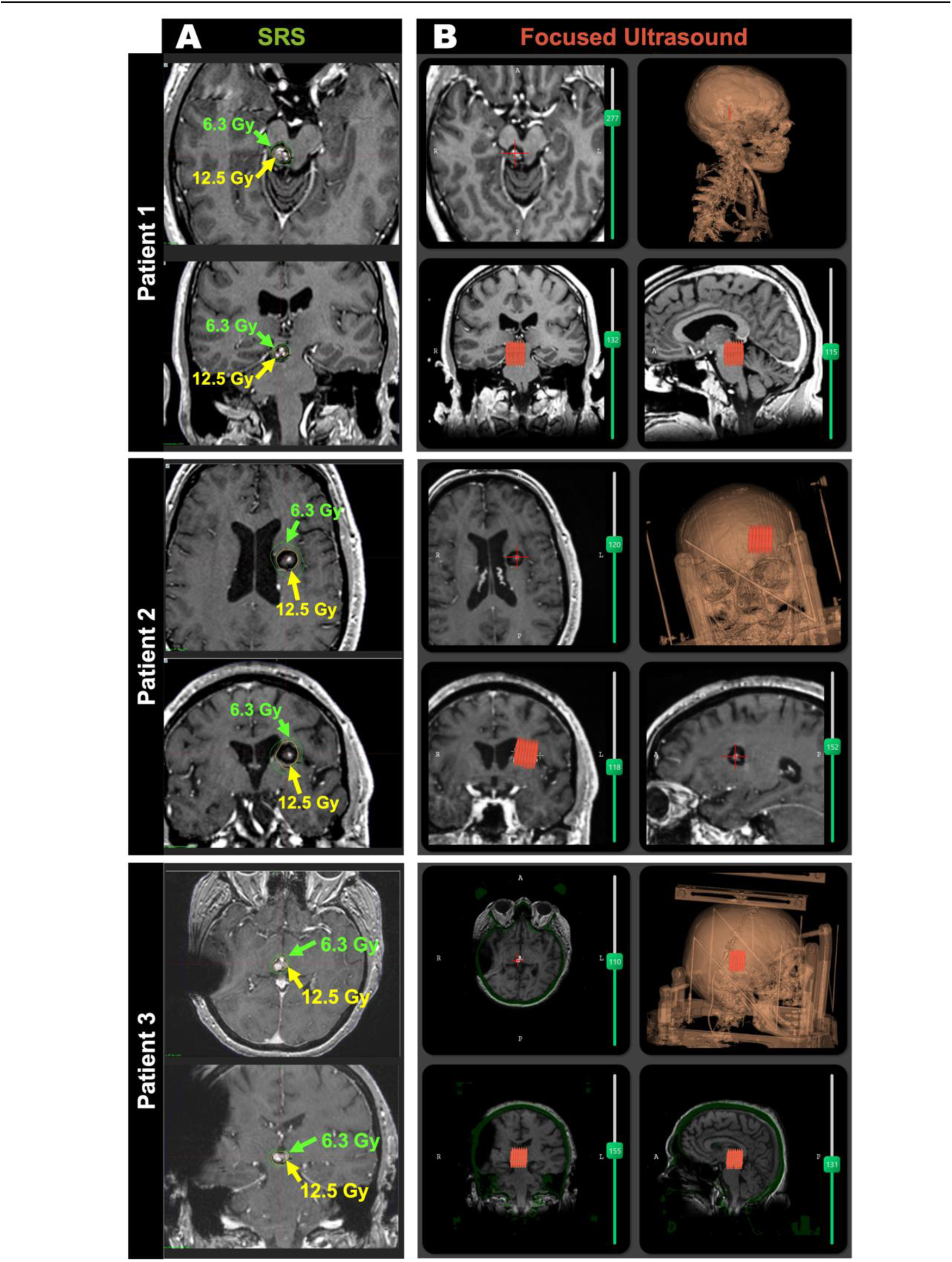
Current clinical FUS systems are equipped to treat CCMs in patients. (A) Stereotactic radiosurgery (SRS) treatment plans for 3 CCM patients with surgically inaccessible lesions. Yellow and green lines are 12.5 Gy and 6.3 Gy isodoses, respectively. (B) Mock FUS BBBO treatment plans using the NaviFUS clinical system software, demonstrating the feasibility of CCM treatment with current clinical FUS systems. Red, grouped focal points denote treatment of CCM with 43 sonication points spanning 2 cm in diameter and 8.65 cm^3^ in volume.

## Discussion

Patients with CCM can sustain incapacitating and even life-threatening neurological symptoms. The only curative treatment option for these patients currently is resection of symptomatic CCMs via invasive neurosurgery, which is associated with a high risk of postoperative morbidities. Further, some CCMs are not surgically accessible^6,35,36^. While SRS may be deployed for patients with inaccessible lesions, SRS can present adverse radiation side effects, induce new CCMs in certain patient populations, and may have limited therapeutic efficacy^37–48^. Concurrently, FUS BBBO is now well-known to exert potentially favorable bioeffects^13,22^. Indeed, we demonstrate here that FUS BBBO can elicit powerful therapeutic effects in a clinically-representative murine model of CCM. Notably, FUS BBBO arrested the growth of 94% of CCMs treated in the study over a 1 month period. Meanwhile, untreated CCMs grew to almost 7 times their initial volume on average across the 3 treatment arms in this same timeframe. Further, mice that received multiple FUS BBBO treatments with fixed PNPs had a significant reduction in the formation of *de-novo* CCMs by 81%. At the cellular level, FUS BBBO reversed the mesenchymal morphology of the CCM vasculature to a more endothelial-like appearance and remodeled the immune landscape of the lesions. As an incisionless therapy with the ability to target eloquent brain locations, FUS BBBO is a disruptive technology that could radically transform how CCMs are treated.

### Characteristics of FUS BBBO in CCM mice

One key consideration in these studies was whether FUS BBBO signatures in *Krit1* mutant mice differ from those in wild-type mice. Since the vasculature associated with CCMs is known to be irregular and dilated^3,49^, the effectiveness of FUS BBBO had the potential to be reduced or otherwise altered. Increased vessel diameters could reduce the interaction between the oscillating microbubbles and vessel walls^50,51^. Moreover, the slow flow rate in the lesion core could reduce the number of microbubbles accumulating within the CCM^49^. Our studies indicate that the pattern of T1 contrast enhancement is localized to the perilesional boundaries of the CCM (**Figure 1B**), which may indicate that the lesion core is not substantially interacting with microbubbles, perhaps due to its grossly enlarged diameter or its slow flow rate. Meanwhile, the perilesional microvasculature displayed marked gadolinium accumulation regardless of moderate vessel diameter dilation compared to normal brain capillaries (**Figure 1D**). Further, our findings suggest that T1 contrast enhancement as well as subharmonic, ultraharmonic, and broadband acoustic signatures of microbubble activity are not significantly different between CCM mice and wild-type mice (**Figure 3**). While the harmonic signatures for PNPs of 0.4 MPa and 0.5 MPa were not significantly different between CCM and wild-type mice, increases in harmonic signatures were seen in CCM mice at 0.6 MPa (**Figure 3E**). This is the only indication that the altered properties of the CCM vasculature, such as vessel diameter, stiffness, and contractility^23–25^, can impact microbubble activity when high enough PNPs are applied. Additionally, since CCMs have a baseline leakiness, it was possible that FUS BBBO would not increase the accumulation of small molecules within the lesion microenvironment. Nevertheless, T1 contrast enhancement from the post-FUS image over the pre-FUS image is indeed apparent for PNPs ranging from 0.3 MPa – 0.6 MPa (**Figure 1D**), indicating that gadolinium accumulation is increased over baseline levels via FUS BBBO. Ultimately, while the pattern of T1 contrast enhancement may be altered in CCM mice, FUS still effectively opens the BBB in the perilesional vasculature of the lesion, and the MRI and acoustic signatures are largely comparable to wild-type mice.

### Acute stability of CCMs exposed to FUS BBBO

The capricious state of these hemorrhage-prone CCMs raised an important concern: would FUS BBBO increase the propensity of CCMs to bleed? The addition of mechanical stress and disruption of already loose endothelial cell tight junctions from oscillating microbubbles had the potential to weaken the stability of CCMs. However, our findings corroborate the safety of FUS BBBO for CCMs. Even susceptibility-weighted images, which have an increased sensitivity to blood products, demonstrated no acute changes in bleeding between the pre- and post-sonication images (**Figure 2D, E**). T2-weighted spin echo sequences, which can accurately represent lesion volume and internal architecture^32^, displayed no acute changes in lesion volume between the pre- and post-sonication images (**Figure 2B, C**). These results also continued for post-sonication images at later timepoints of up to one month, indicating that FUS BBBO is safe for CCMs both acutely and chronically. Meanwhile, our results did indicate that edema and hemosiderin deposits can be seen in lesion-free brain tissue in *both* wild-type and CCM mice when using PNPs greater than 0.4 MPa (**Figure 4**). This finding further supports the use of PCD-modulated PNP feedback systems that have been widely adopted in clinical trials to ensure the safety of FUS BBBO treatments (**Figure 5**)^26,28–31,52^.

### FUS BBBO provides a therapeutic effect for CCMs and familial forms of the disease

After establishing that FUS BBBO was safe, we questioned whether it could be therapeutic for CCMs. From analysis of longitudinal MR images, we show that FUS BBBO is capable of both fully arresting the growth of pre-existing CCMs (**Figure 6**) and preventing *de novo* CCM formation (**Figure 7**). The ability to slow and even reverse the growth of CCMs could have far-reaching implications for CCM therapy. The pathological trajectory of many CCMs remains uncertain to clinicians^9–11^, so patients must choose between neurosurgery, with its associated risks^6,7^, or inaction. FUS BBBO could provide a non-invasive alternative to enable the stabilization of the lesion without the risks associated with surgery or the lack of intervention.

Further, this approach could be revolutionary for patients with the familial form of the disease.

Familial CCM patients have multiple lesions, of which several can often arise in locations that are inoperable or are associated with a very high risk for post-operative morbidities^6,9^. FUS BBBO could be used to stabilize multiple CCMs within a single treatment session, including those in eloquent locations, while simultaneously protecting those sonicated areas from future CCMs. FUS BBBO could help make an impossible choice for CCM patients and parents of CCM patients more manageable.

### Potential mechanisms for the protective effect of FUS BBBO in CCM

The ability of FUS BBBO to exert powerful therapeutic effects for CCMs was surprising; however, this is not the first disease indication wherein FUS BBBO has been shown to be protective. FUS BBBO—in the absence of drug delivery—has also exhibited a therapeutic effect for Alzheimer’s disease^16–22^. The exact mechanism of action in Alzheimer’s disease remains unclear, though many studies have investigated the potential mechanisms behind its benefit^18–21,53^.

In this study, our extensive histological analysis lends some insight into how FUS BBBO may benefit CCMs. At 1 month post-FUS BBBO, when growth control is evident for all FUS BBBO regimens, the mesenchymal morphology of the mutated CCM vasculature was restored to an endothelial morphology in sonicated lesions (**Figure 8A-B**). Thus, FUS BBBO appears to reverse the endothelial-to-mesenchymal transition that is common in CCMs.

Because FUS BBBO increases phagocytosis in other disease contexts^33,34^, another hypothesis for CCM stabilization was that FUS BBBO-exposed microglia and macrophages would become activated and phagocytose erythrocytes. However, our data are not consistent with this putative mechanism of lesion control. Instead, we found that the co-localization of Iba1+ microglia/macrophages with erythrocytes was actually decreased at 7 days post-FUS BBBO (**Figure 8F, H**). Beyond microglia and macrophages, numerous studies indicate that FUS BBBO increases immune cell infiltration in a variety of disease states^13,34,54–57^. Consistent with these studies, we confirmed that FUS BBBO increases overall immune cell (CD45+) infiltration in CCMs (**Figure S5**), signifying an altered immune landscape as a potential mechanism for CCM stabilization. Ultimately, several mechanisms may underlie the protective role of FUS BBBO for CCM.

### The potential of FUS BBBO to synergize with pharmacological treatments

To date, no pharmacological agent has been approved for the treatment of CCM, yet a few drugs have entered clinical trials (propranolol: NCT03589014, REC-994: NCT05085561, simvastatin: NCT01764451, and atorvastatin: NCT02603328). Additionally, many drugs for CCM are being examined in the preclinical stage^1^. These drug candidates have the potential to seamlessly integrate with the FUS BBBO approach used in this study, especially since surgically inaccessible CCMs in eloquent regions are accessible for FUS BBBO using current clinical FUS systems (**Figure 9**). Therapeutic agents can be injected alongside FUS BBBO and benefit from the enhanced permeability as a way to shift the systemic dose to be more localized to the CCM. This would be reflected as an increase in the therapeutic index, which could be leveraged to reduce the amount of drug needed and help mitigate potential drug side effects. Moreover, FUS BBBO could also have the potential to unlock whole new classes of drug candidates. Larger molecular weight biologics, like antibodies and gene therapies, would have a greater potential to accumulate in the CCM microenvironment with the aid of increased permeability via FUS BBBO^13,14,58^. Indeed, the vast majority of the drug candidates being studied for CCM currently are small molecules^1^. Ultimately, the innate protective effect of FUS BBBO for CCM and the countless drug candidates that could integrate with the enhanced delivery of this approach provides an immeasurable potential to vastly expand the therapeutic options and to transform the treatment paradigm for CCM.

## Methods

### Animals

All animal experiments were approved by the University of Virginia Animal Care and Use Committee. Mice were housed under standard laboratory conditions (22°C and 12h/12h light/dark cycle). The generation of the CCM murine models (*Pdgfb-CreERT2;Krit1^fl/null^*or *Cdh5-CreERT2;Krit1^fl/null^*) that were used in these studies has been described previously^32^. Briefly, *Pdgfb-CreERT2* or *Cdh5-CreERT2* mice were crossed with *Krit1*^fl/null^ male or females. On postnatal day 5, *Krit1* gene ablation was induced with an injection of tamoxifen (subcutaneous; 50uL at 2mg/mL in corn oil). Genotypes were confirmed using Transnetyx (Cordova, TN). Wild-type mice in this study were on the same background strain as the CCM model (C57BL/6; Charles River). All mice were treated between 9 weeks and 13 weeks of age. Mouse sex, litter, age, and treatment assignment are listed in detail in **Table S1**.

### MR Imaging

MR imaging was performed using either a 7T Bruker/Siemens ClinScan or a 9.4T Bruker BioSpec small animal MRI scanner. T2-weighted spin echo images were acquired at 7T with the Siemens 3D T2-SPACE sequence (repetition time of 3000 ms, echo time of 80 ms, pixel size of 125 μm x 125 μm x 100 μm, 2 averages, and 20 min acquisition time) or at 9.4T with the Bruker 3D T2-TurboRARE sequence (repetition time of 2000 ms, echo time of 55 ms, turbo factor of 18, pixel size of 125 μm x 125 μm x 125 μm, 1 average, and 30 min acquisition time). Susceptibility-weighted images were acquired only at 7T (repetition time of 18 ms, echo time of 10 ms, pixel size of 130 μm x 130 μm x 130 μm, 2 averages, and 15 min acquisition time). T1-weighted spin echo images were acquired at 9.4T with the Bruker 2D T1-RARE sequence (repetition time of 1500 ms, echo time of 6 ms, pixel size of 156 μm x 156 μm x 350 μm, 1 average, and 3 min acquisition time). All imaging was performed under isoflurane anesthesia, and body temperature was maintained with a heated, circulating water bed.

### Selection of CCMs for Sonication

Following baseline MR image acquisition, images were reviewed to assess appropriate CCMs for sonication. CCMs located within the left or right caudoputamen, corpus callosum, or cerebral cortex were eligible for targeting. The average sonicated and non-sonicated (contralateral control) CCM volume prior to FUS application was 0.039 mm^3^ for both conditions in the longitudinal studies. Prior to safety evaluation measurements and analysis, sonications were confined to single CCMs without neighboring CCMs located dorsally or ventrally that would be within the focal zone. Following the initial safety evaluation, multiple CCMs were eligible for sonication if they were within the same focal volume.

### FUS BBBO

FUS BBBO was performed with the RK-300 small bore FUS device (FUS Instruments, Toronto, CA). Heads of mice were shaved and depilated prior to supine placement and coupling to the transducer with degassed ultrasound gel. BBBO was performed with a 1.13 MHz single-element transducer using a 10 ms burst length over a 2000 ms period for 60 total sonications during a 2-min sonication duration. Fixed PNP application was performed using the “Burst” mode on the FUS Instruments software. PCD-modulated PNP was performed using the “Blood-brain Barrier” mode of the FUS Instruments software. Parameters used for this feedback control system included a starting pressure of 0.2 MPa, pressure increment of 0.05 MPa, maximum pressure of 0.4 MPa, 20 sonication baselines without microbubbles, AUC bandwidth of 500 Hz, AUC threshold of 10 standard deviations, pressure drop of 0.95, and frequency selection of the subharmonic, first ultraharmonic, and second ultraharmonic. Gadolinium contrast agent (Multihance) was injected as a bolus intravenously with a dose of 0.01 mmol diluted in saline at a molarity of 0.2 mmol/mL prior to T1-RARE image acquisition. Albumin-shelled microbubbles were made in-house as previously described^59^ and intravenously injected as a bolus dose of 10^5^ microbubbles per gram body weight. Distribution of microbubble diameter and concentration was acquired with a Coulter counter (Multisizer 3; Beckman Coulter, Fullerton, California) prior to sonication. High resolution T2-weighted images and T1-RARE images were used to guide FUS targeting to the pre-selected CCM. A single sonication target was used in all experiments, except in the case of PCD-modulated PNPs, in which two sonication targets were used. Mice receiving the repeat FUS BBBO regimens had all sonications staged 3 days apart with the same anatomical location targeted each time.

### Acoustic Signatures from Passive Cavitation Detection

Acoustic emissions were detected with a fiber-optic hydrophone (Precision Acoustics, Dorset, UK) of 10 mm diameter and 15 mm aperture center-mounted within the ultrasound transducer. Emissions data was processed and spectrograms were generated with a custom MATLAB script. The area under the curve of the acoustic emissions at the subharmonic (0.5f) and ultra-harmonics (1.5f, 2.5f) were calculated after applying a 300 Hz bandwidth filter. Broadband emissions were evaluated by summing acoustic emissions following the removal of all emissions at the fundamental frequency (f), harmonics (2f, 3f, 4f), subharmonic (0.5f), and ultra-harmonics (1.5f, 2.5f, 3.5f).

### T1 Contrast Enhancement Analysis

Gadolinium accumulation following FUS BBBO was evaluated using the enhancement of T1 contrast in T1-RARE images. In a DICOM viewer (Horos Project, Geneva, Switzerland), an ROI was drawn around the boundaries of the enhanced (hyperintense) region on the image slice containing the targeted lesion. The ROI was then copied onto the pre-sonication T1-RARE image on the same slice. For wild-type mice, ROIs were drawn around the boundaries of the enhanced (hyperintense) region in similar ventral/dorsal slice depths as CCM mice. Mean grayscale intensity for each ROI was recorded, and fold change in grayscale intensity from the post-image to the pre-image was calculated. This process was repeated for all sonicated mice across each PNP.

### Brain Tissue Edema and Hemosiderin Deposition Analysis

Edema and hemosiderin deposition in lesion-free brain tissue following FUS BBBO were evaluated in 3D Slicer using the high resolution T2-weighted spin echo MR images. MR images were initially segmented by the brain tissue boundaries to generate a mask of the brain. Bias field correction was then applied with the N4ITK MRI Bias Field Correction tool in 3D Slicer to correct for inhomogeneities in signal intensity across the brain due to mouse rotation relative to the MR surface coil. Mean grayscale intensity was then recorded within ROIs of equal volume in lesion-free brain tissue for both non-sonicated (contralateral) and sonicated (ipsilateral) hemispheres on the same dorsal slice. Healthy brain tissue would have an ipsilateral-to-contralateral grayscale ratio near 1. Edema would produce a ratio greater than 1, while hemosiderin would produce a ratio less than 1.

### CCM Growth Analysis

CCM volume prior to, and longitudinally following, FUS BBBO was evaluated in Horos using the high resolution T2-weighted spin echo MR images. For each timepoint, an ROI was manually drawn around the sonicated CCM in each slice it was present. The Horos “Compute Volume” tool was then used to calculate the three-dimensional volume of the CCM across imaging timepoints. In the same mice, ROIs were also drawn around non-sonicated CCMs (i.e. control CCMs) that had similar volumes and anatomical locations as sonicated lesions. CCM mice with enlarged ventricles, a rare but potential co-morbidity of this model, at the one-month timepoint were removed from this analysis.

### New Lesion Formation Analysis

Formation of new CCMs was assessed by calculating the change in lesion number from the baseline pre-FUS to the one-month post-FUS high resolution T2-weighted spin echo MR images. For both timepoints, an ROI was first drawn around the T1 contrast enhanced boundaries within the T1-RARE images taken following FUS BBBO, extending from the most dorsal to most ventral slices of the brain and focal column. These ROIs were then copied onto the T2-weighted spin echo images and adjusted to match the same anatomical positioning. These ROIs were then copied to the contralateral brain region and adjusted to mirror the same anatomical positioning. CCMs within the ROIs were then manually counted and recorded for both timepoints and for both the ipsilateral ROI and the contralateral ROI. The baseline CCM number was subtracted from the one-month CCM number for both the ipsilateral ROI volume and the contralateral ROI volume in each mouse to produce the number of new CCMs formed in each ROI volume during the one-month time period. CCM mice with enlarged ventricles at the one-month timepoint were removed from this analysis.

### Immunohistochemistry

Mice were perfused with phosphate-buffered saline (PBS) and 4% paraformaldehyde, and after harvesting, brains were fixed overnight in 4% paraformaldehyde and dehydrated in 30% sucrose solution for 24 h. Brains were then embedded in Optimal Cutting Temperature Compound (TissueTek) for cryosectioning at 30-µm thickness. Sections were incubated in blocking solution (1% bovine serum albumin, 2% normal donkey serum, and 0.1% Triton X-100, and 0.05% Tween-20 in PBS) for 2 h at RT. Brain sections were then incubated with goat anti-CD31 (1:20, R&D Systems, AF3628), rat anti-GFAP-Alexa Fluor 488 (1:50, eBioscience, 53-9792-82), rat anti-Ki67-Alexa Fluor 660 (1:100, ThermoFisher, 50-5698-82), rabbit anti-Iba1 (1:400, FujiFilm Wako, 019-19741), rat anti-CD68-Alexa Fluor 700 (1:50, BioRad, MCA1957A700), rat anti-Ter119-Super Bright 436 (1:100, ThermoFisher, 62-5921-82), and goat anti-CD45 (1:200, R&D Systems, AF114) diluted in the blocking solution overnight at 4°C. After three 5-min washes in PBS with 0.5% Tween-20, the sections were incubated with donkey anti-goat-Alexa Fluor 647 (1:500, Invitrogen A21447), donkey anti-rabbit-Alexa Fluor 405 (1:1000, ThermoFisher, A48258), donkey anti-rabbit-Alexa Fluor 488 (1:1000, Abcam, ab150073), and donkey anti-goat-Alexa Fluor 405 (1:1000, Abcam, ab175664) and diluted in the blocking solution for 2 h at RT. Sections were imaged with a Leica Stellaris 5 confocal microscope (Leica Microsystems). Images were processed with Fiji/ImageJ.

### Analysis of Immunofluorescent Images

Images were collected as a z-stack of 1-µm step size at either 20x or 63x magnification. For 20x images, tiled images were collected to cover the perilesional and intralesional space of sonicated and non-sonicated CCMs. For 63x images, non-tiled images were acquired along the perilesional and intralesional boundary of sonicated and non-sonicated lesions. Maximum intensity projections were produced in Fiji/ImageJ. Quantification of cell markers, morphology, and colocalization was conducted in HALO using the Object Colocalization and Highplex modules.

### Statistical Analysis

All results are reported as mean ± standard error of the mean (SEM). The “n” values per group are made evident either by individual data points shown or statement of “n” value in figure, figure legend, and/or manuscript text. Statistical significance was assessed at p < 0.05 for all experiments. Linear mixed effect models were conducted and analyzed with the lme4 package (version 1.1.34) and the emmeans package (version 1.8.9) in R Studio. All other statistical tests were performed using GraphPad Prism 9 (San Diego, USA). Statistical tests, models, and p-values are listed in detail for all manuscript figures in **Table S2**.

## Author Contributions

DGF and RJP conceptualized the study. DGF conducted the FUS BBBO experiments with the aid of CMG, VRB, ACD, MRH, and TC in animal preparation and MRI acquisition. MRI sequences were optimized by GWM. Longitudinal MRI data was acquired by DGF and analyzed by DGF and IMS. KAS, PT, TC, and DGF generated experimental animals. DGF and KAS performed immunostaining and confocal imaging with technical guidance from JDS and JRL. JPS and DS provided SRS clinical treatment plans, and DM provided FUS BBBO clinical treatment plans. DGF designed the figures and wrote the manuscript. JDS, ACD, JSP, JRL, PT, GWM, and RJP edited the manuscript. All authors approved the manuscript.

## Acknowledgements

Supported by funding from NIH R01CA279134, R01EB030409, R01EB030744, and R21NS118278 to RJP; NIH R21NS116431 and grants from Focused Ultrasound Foundation, Be Brave for Life Foundation, and Alliance to Cure Cavernous Malformation to PT; NIH R01CA226899 to GWM; and AHA 830909 to DGF. We thank Dr. Kevin Whitehead for kindly providing the mouse strains used in this study. We are also grateful to the University of Virginia Molecular Imaging Core and Jeremy Gatesman of the University of Virginia Center for Comparative Medicine for assistance with MRI imaging and catheterization procedures, respectively. Additionally, we extend our gratitude to Marieke Jones of the University of Virginia School of Medicine for her biostatistical guidance.

