## Supplemental Material for "Focused Ultrasound Blood-Brain Barrier Opening Arrests the Growth and Formation of Cerebral Cavernous Malformations"

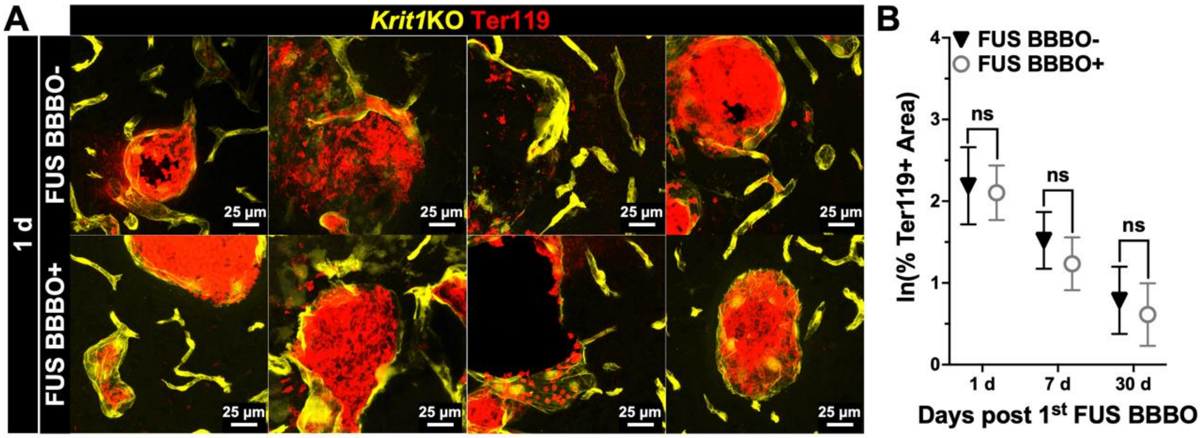


**Figure S1.** FUS BBBO does not exacerbate CCM hemorrhage. (A) Immunofluorescent images of non-sonicated and sonicated CCMs at 1 d post FUS BBBO with staining for mutated vasculature *(Krit1*KO) and erythrocytes (Ter119)*.* (B) Graph of the percent of erythrocyte positive area at 1 d, 7 d, and 30 d post FUS BBBO for non-sonicated and sonicated CCMs, indicating no significant differences in erythrocyte coverage at any time point post FUS BBBO. Linear mixed effect model and pairwise comparison with Tukey's adjustment.

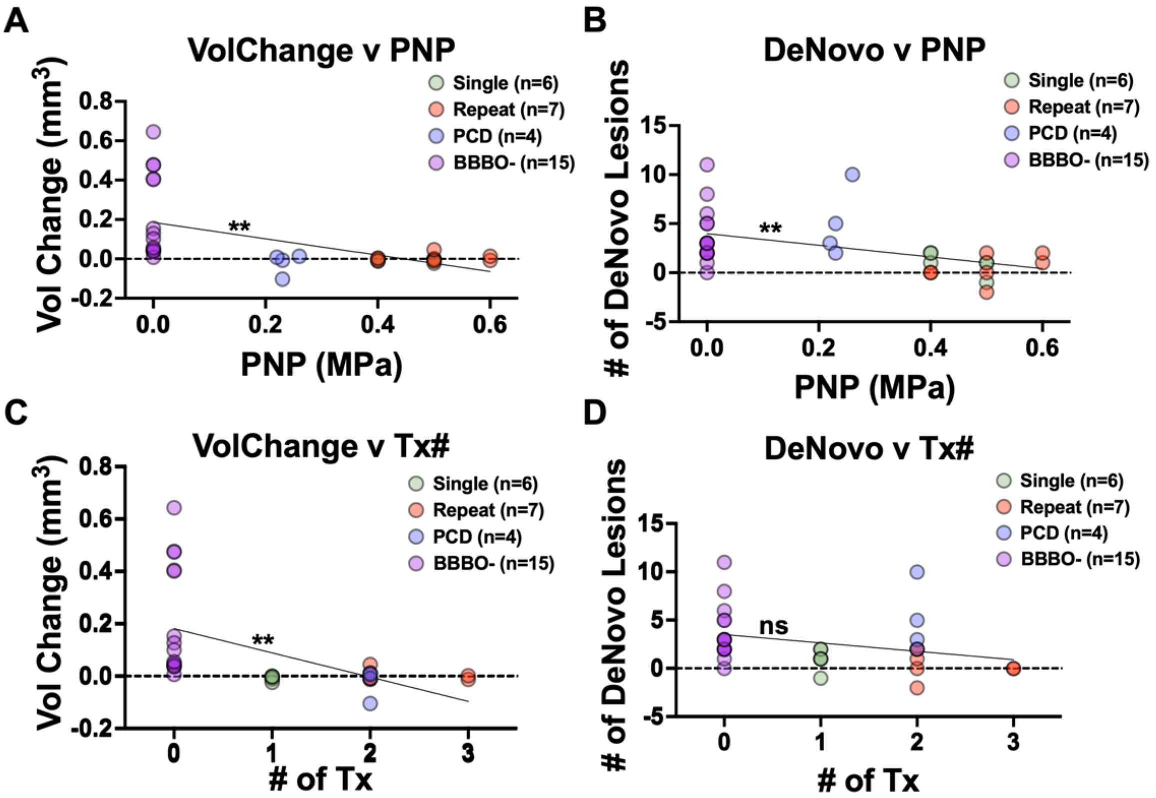


**Figure S2.** Correlations of FUS BBBO PNP and treatment number on CCM volume change and formation. (A) Plot of lesion volume change and PNP for all treatment conditions. Linear regression indicates that volume change and PNP are inversely correlated, $p = 0.0017$. (B) Plot of *de novo* CCM formation and PNP for all treatment conditions. Linear regression indicates that *de novo* CCM formation and PNP are inversely correlated, $p = 0.0064$. (C) Plot of lesion volume change and number of FUS applications (i.e. Tx#) for all treatment conditions. Linear regression indicates that volume change and Tx# are inversely correlated, $p = 0.0021$. (D) Plot of *de novo* CCM formation and number of FUS applications (i.e. Tx#) for all treatment conditions. Linear regression indicates that *de novo* CCM formation and Tx# are not inversely correlated, $p = 0.0914$.

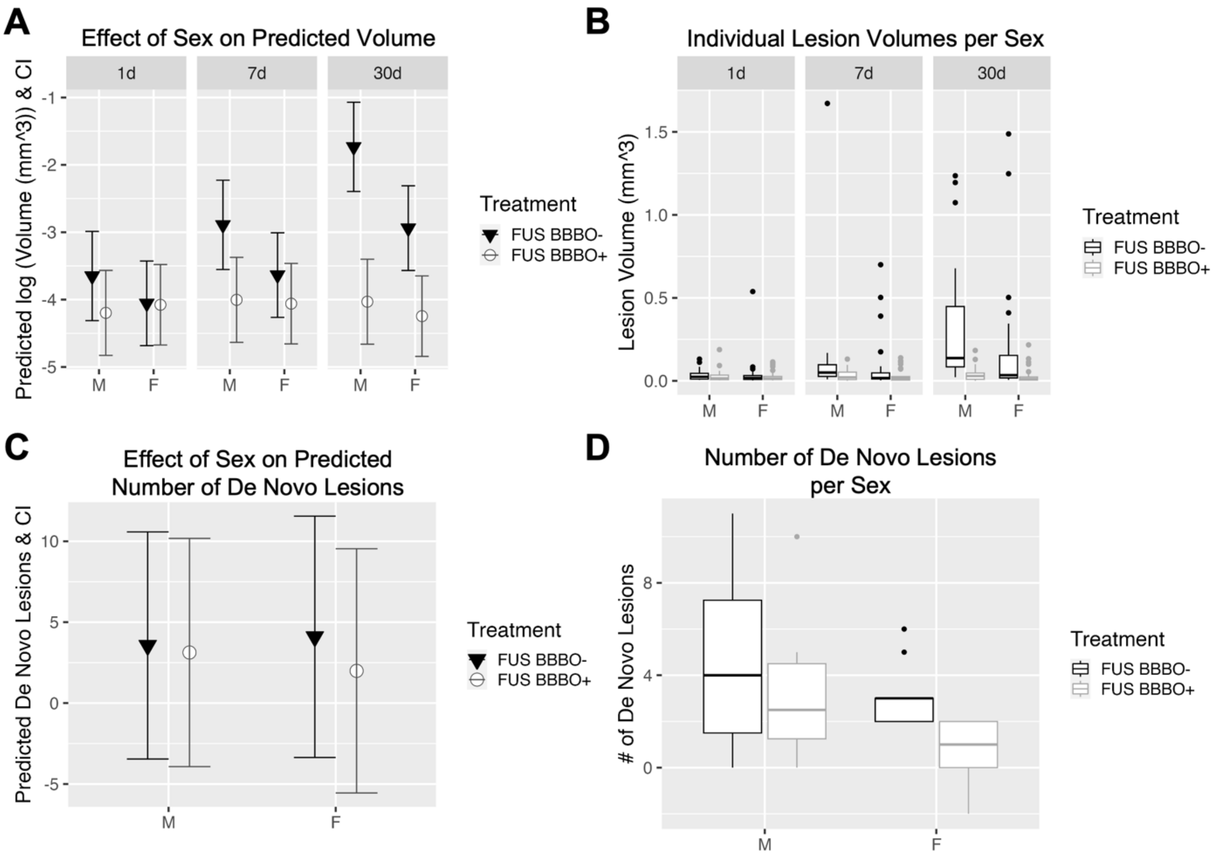


**Figure S3.** Effect of sex on the therapeutic outcomes of FUS BBBO for CCM. (A) Summary plots comparing the natural log transform of CCM volume between sonicated CCMs and non-sonicated CCMs over time disaggregated by sex. At 1 month, CCMs in male mice were larger than those in female mice, regardless of FUS BBBO treatment (Linear mixed effect model: Table S2). Sex did not significantly alter the ability of FUS BBBO to control CCM growth (Linear mixed effect model: Table S2). (B) Plots of CCM volume for individual CCMs for each treatment group over time disaggregated by sex. (C) Summary plots comparing the number of *de novo* CCMs between sonicated CCMs and non-sonicated CCMs disaggregated by sex. Sex did not significantly alter the ability of FUS BBBO to control CCM formation (Linear mixed effect model: Table S2). (D) Boxplots comparing the number of *de novo* CCMs for each treatment group disaggregated by sex.

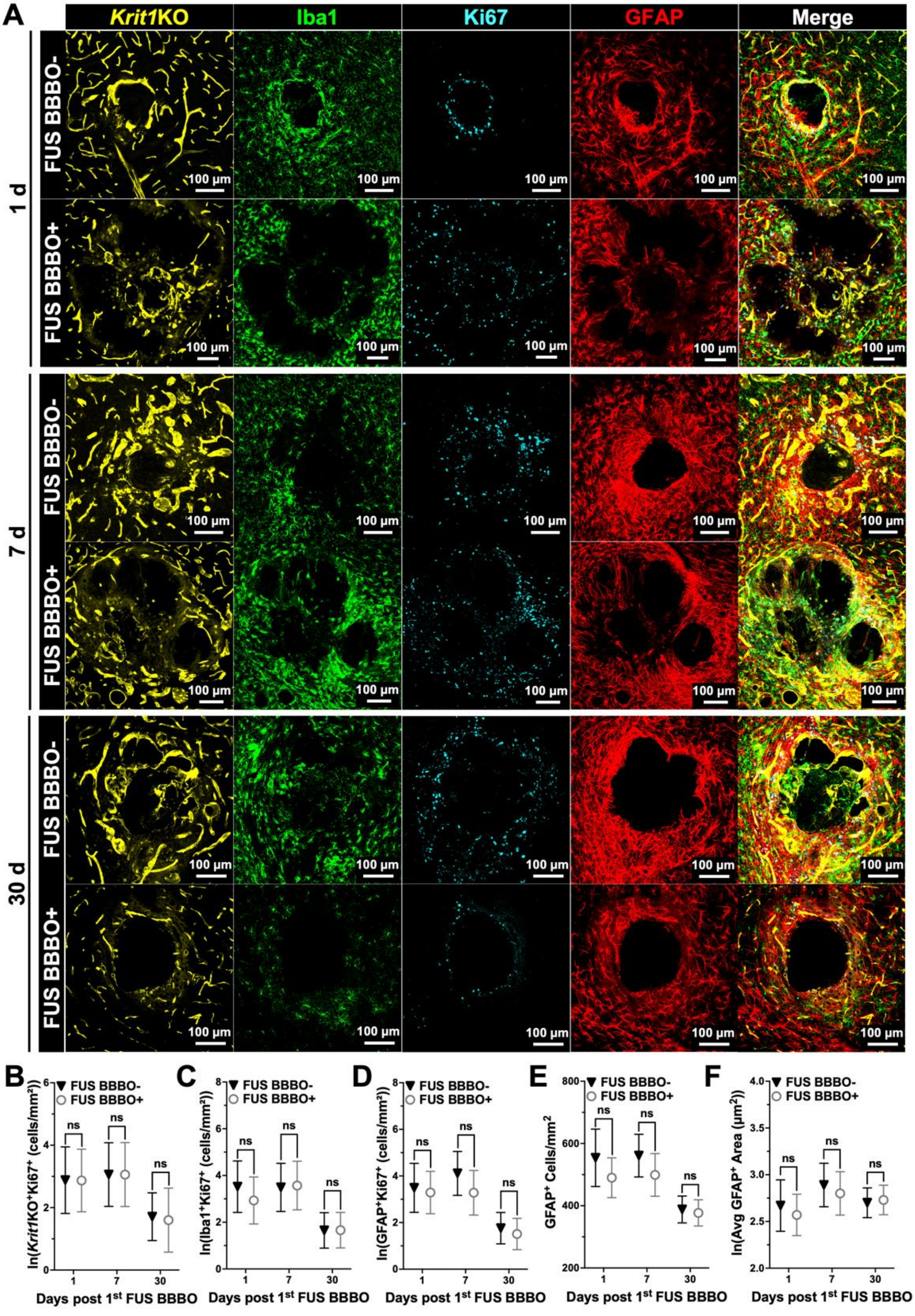
A
Krit1KO
Iba1
Ki67
GFAP
Merge
1 d
7 d
30 d
FUS BBBO-
FUS BBBO+
100 µm
B
C
D
E
F
FUS BBBO-
FUS BBBO+
ns
ln(Krit1KO⁺Ki67⁺ (cells/mm²))
ln(Iba1⁺Ki67⁺ (cells/mm²))
ln(GFAP⁺Ki67⁺ (cells/mm²))
GFAP⁺ Cells/mm²
ln(Avg GFAP⁺ Area (µm²))
Days post 1st FUS BBBO

56

**Figure S4.** FUS BBBO does not change proliferation of mutated endothelial cells or lesion-associated microglia/macrophages and astrocytes. (A) Immunofluorescent images of non-sonicated and sonicated CCMs at 1 d, 7d, and 30 d post FUS BBBO with staining for mutated vasculature *(Krit1*KO), microglia/macrophages (Iba1), proliferation (Ki67), and astrocytes (GFAP). (B-D) Graphs of the natural log of Ki67+ mutated endothelial cells, Ki67+ microglia/macrophages, and Ki67+ astrocytes, respectively, at 1 d, 7 d, and 30 d post FUS BBBO for non-sonicated and sonicated CCMs, indicating no significant differences in proliferation of these cells at any time point post FUS BBBO. Linear mixed effect model and pairwise comparison with Tukey's adjustment. (E-F) Graph of astrocyte density and the natural log of astrocyte area at 1 d, 7 d, and 30 d post FUS BBBO for non-sonicated and sonicated CCMs, demonstrating no significant differences in astrocyte number or size following FUS BBBO. Linear mixed effect model and pairwise comparison with Tukey's adjustment.

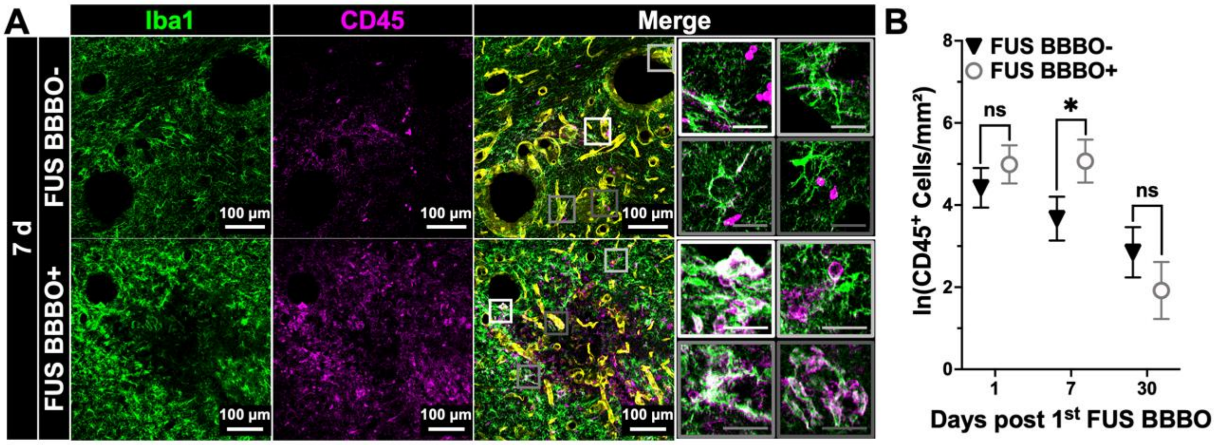


**Figure S5.** FUS BBBO increases immune cell infiltration into CCMs 7 d post sonication. (A) Immunofluorescent images of non-sonicated and sonicated CCMs at 7 d post FUS BBBO with staining for microglia/macrophages (Iba1), immune cells (CD45), and mutated vasculature (*Krit1*KO). Insets display magnified views of the corresponding 20x image, revealing monocytes in the lumens of CCMs, microglia/macrophage processes extending to immune cells, and CD45+ macrophages lining mutated vessels. *(*B) Graph of the natural log of CD45+ immune cell density at 1 d, 7 d, and 30 d post FUS BBBO for non-sonicated and sonicated CCMs, demonstrating a marked increase in immune cell infiltration at 7 d. $p = 0.0494$, linear mixed effect model and pairwise comparison with Tukey's adjustment.

**Table S1:** Mouse and treatment assignment characteristics

| Mouse ID | Tx Cohort | PNP (or AVG PNP) | Litter # | Sex | Age at Tx (wk) |
|---|---|---|---|---|---|
| 32 | Repeat | 0.5 | 1 | F | 10 |
| 33 | Repeat | 0.6 | 1 | F | 10 |
| 37 | Repeat | 0.4 | 2 | M | 10 |
| 38 | Repeat | 0.6 | 2 | M | 10 |
| 103 | Repeat | 0.5 | 3 | F | 9 |
| 104 | Repeat | 0.4 | 3 | F | 9 |
| 106 | Repeat | 0.4 | 3 | F | 9 |
| 81 | Repeat | 0.5 | 4 | F | 10 |
| 83 | Repeat | 0.6 | 4 | M | 10 |
| 903 | Single | 0.5 | 5 | F | 9 |
| 904 | Single | 0.4 | 5 | F | 9 |
| 74 | Single | 0.5 | 6 | F | 9 |
| 75 | Single | 0.5 | 6 | F | 9 |
| 76 | Single | 0.4 | 6 | F | 9 |
| 79 | Single | 0.4 | 6 | M | 9 |
| 841 | PCD | 0.26 | 7 | M | 13 |
| 831 | PCD | 0.23 | 8 | M | 13 |
| 306 | PCD | 0.22 | 9 | M | 11 |
| 311 | PCD | 0.23 | 9 | F | 11 |

**Table S2:** Statistical tests and p-values for manuscript figures

| | |
|---|---|
| **Figure 1C: One-way ANOVA with Dunnett's Multiple Comparisons Test** | |
| Comparison | p-value |
| Pre vs. 0.2 | 0.9986 |
| Pre vs. 0.3 | 0.0054 |
| Pre vs. 0.4 | <0.0001 |
| Pre vs. 0.5 | <0.0001 |
| Pre vs. 0.6 | <0.0001 |
| **Figure 2B: Wilcoxon Matched-Pairs Signed Rank Test** | |
| Comparison | p-value |
| Pre-FUS vs. Post-FUS | 0.4143 |
| **Figure 2D: Wilcoxon Matched-Pairs Signed Rank Test** | |
| Comparison | p-value |
| Pre-FUS vs. Post-FUS | 0.3396 |
| **Figure 3B: Two-way ANOVA with Šidák's Multiple Comparisons Test** | |
| Comparison | p-value |
| 0.4 MPa: WT vs. CCM | 0.9238 |
| 0.5 MPa: WT vs. CCM | 0.9998 |
| 0.6 MPa: WT vs. CCM | 0.9590 |
| **Figure 3D: Two-way ANOVA with Šidák's Multiple Comparisons Test** | |
| Comparison | p-value |
| Subharmonic: 0.4 MPa: WT vs. CCM | 0.8640 |
| Subharmonic: 0.5 MPa: WT vs. CCM | 0.9864 |
| Subharmonic: 0.6 MPa: WT vs. CCM | 0.9573 |
| 1st Ultraharmonic: 0.4 MPa: WT vs. CCM | 0.9058 |
| 1st Ultraharmonic: 0.5 MPa: WT vs. CCM | 0.9934 |
| 1st Ultraharmonic: 0.6 MPa: WT vs. CCM | 0.8039 |
| Broadband: 0.4 MPa: WT vs. CCM | 0.8457 |
| Broadband: 0.5 MPa: WT vs. CCM | 0.9900 |
| Broadband: 0.6 MPa: WT vs. CCM | 0.4019 |
| **Figure 3E: Two-way ANOVA with Šidák's Multiple Comparisons Test** | |
| Comparison | p-value |
| 2nd Harmonic: 0.4 MPa: WT vs. CCM | >0.9999 |
| 2nd Harmonic: 0.5 MPa: WT vs. CCM | 0.9998 |
| 2nd Harmonic: 0.6 MPa: WT vs. CCM | <0.0001 |
| 3rd Harmonic: 0.4 MPa: WT vs. CCM | >0.9999 |
| 3rd Harmonic: 0.5 MPa: WT vs. CCM | >0.9999 |
| 3rd Harmonic: 0.6 MPa: WT vs. CCM | 0.0006 |
| 4th Harmonic: 0.4 MPa: WT vs. CCM | 0.9999 |
| 4th Harmonic: 0.5 MPa: WT vs. CCM | 0.7607 |
| 4th Harmonic: 0.6 MPa: WT vs. CCM | <0.0001 |
| **Figure 4B: 2nd Order Polynomial Regression Comparison of Fits** | |
| Null hypothesis | One curve for all data sets |
| Alternative hypothesis | Different curve for each data set |

| | |
|---|---|
| p-value | 0.0473 |
| **Figure 4C: 2nd Order Polynomial Regression Comparison of Fits** | |
| Null hypothesis | One curve for all data sets |
| Alternative hypothesis | Different curve for each data set |
| p-value | 0.7734 |
| **Figure 4E: Two-way ANOVA with Holm-Šidák's Multiple Comparisons Test** | |
| Comparison | p-value |
| Single: Edema: 0.4 MPa: WT vs. CCM | 0.1368 |
| Single: Edema: 0.5 MPa: WT vs. CCM | 0.1368 |
| Single: Hemosiderin: 0.4 MPa: WT vs. CCM | 0.5386 |
| Single: Hemosiderin: 0.5 MPa: WT vs. CCM | 0.5386 |
| Repeat: Edema: 0.4 MPa: WT vs. CCM | 0.7316 |
| Repeat: Edema: 0.5 MPa: WT vs. CCM | 0.9667 |
| Repeat: Edema: 0.6 MPa: WT vs. CCM | 0.0923 |
| Repeat: Hemosiderin: 0.4 MPa: WT vs. CCM | 0.9965 |
| Repeat: Hemosiderin: 0.5 MPa: WT vs. CCM | 0.5754 |
| Repeat: Hemosiderin: 0.6 MPa: WT vs. CCM | 0.9965 |
| **Figure 5C: Wilcoxon Matched-Pairs Signed Rank Test** | |
| Comparison | p-value |
| Pre-FUS vs. Post-FUS | 0.0156 |
| **Figure 5D: One-way ANOVA with Dunnett's Multiple Comparisons Test** | |
| Comparison | p-value |
| PCD vs. 0.2 | <0.0001 |
| PCD vs. 0.3 | 0.0018 |
| PCD vs. 0.4 | 0.0368 |
| PCD vs. 0.5 | 0.2864 |
| PCD vs. 0.6 | 0.9918 |
| **Figure 5F: Two-way ANOVA with Šidák's Multiple Comparisons Test** | |
| Comparison | p-value |
| Subharmonic: PCD vs. 0.4 | 0.9993 |
| Subharmonic: PCD vs. 0.5 | >0.9999 |
| Subharmonic: PCD vs. 0.6 | 0.3381 |
| Subharmonic: 0.4 vs. 0.5 | 0.9995 |
| Subharmonic: 0.4 vs. 0.6 | 0.5210 |
| Subharmonic: 0.5 vs. 0.6 | 0.3475 |
| Broadband: PCD vs. 0.4 | 0.3359 |
| Broadband: PCD vs. 0.5 | 0.5648 |
| Broadband: PCD vs. 0.6 | 0.6860 |
| Broadband: 0.4 vs. 0.5 | 0.9979 |
| Broadband: 0.4 vs. 0.6 | 0.0538 |

| | |
|---|---|
| Broadband: 0.5 vs. 0.6 | 0.0954 |
| 2nd Harmonic: PCD vs. 0.4 | >0.9999 |
| 2nd Harmonic: PCD vs. 0.5 | >0.9999 |
| 2nd Harmonic: PCD vs. 0.6 | 0.0032 |
| 2nd Harmonic: 0.4 vs. 0.5 | >0.9999 |
| 2nd Harmonic: 0.4 vs. 0.6 | 0.0027 |
| 2nd Harmonic: 0.5 vs. 0.6 | 0.0031 |
| **Figure 5I: Linear Regression** | |
| Edema: Is slope significantly non-zero? | |
| F | 0.05758 |
| DFn, DFd | 1, 17 |
| P value | 0.8132 |
| Equation | Y = 0.03146*X + 0.9876 |
| Hemosiderin: Is slope significantly non-zero? | |
| F | 6.897 |
| DFn, DFd | 1, 17 |
| P value | 0.0177 |
| Equation | Y = -0.3262*X + 1.066 |

| | | | |
|---|---|---|---|
| **Figure 6B: Linear Mixed Effect Model by Restricted Maximum Likelihood** | | | |
| lnVolume ~ Treatment * Time + Starting Volume + (1 \| Mouse) + (1 \| Lesion) | | | |
| Random Effects: | | | |
| <u>Groups</u> | <u>Name</u> | <u>Variance</u> | <u>Std. Dev.</u> |
| Lesion ID | (Intercept) | 0.1452 | 0.3811 |
| Mouse ID | (Intercept) | 0 | 0 |
| Residual | | 0.2248 | 0.4742 |
| Fixed Effects: | | | |
| <u>Groups</u> | <u>Estimate</u> | <u>Std. Error</u> | <u>p-value</u> |
| Intercept | -5.0212 | 0.2103 | 2.00E-16 |
| FUS BBBO+ | -0.156 | 0.2569 | 0.546599 |
| 7d | 0.2827 | 0.2121 | 0.189679 |
| 30d | 1.0516 | 0.2121 | 1.22E-05 |
| Starting Volume | 31.1736 | 5.2957 | 9.30E-06 |
| FUS BBBO+:7d | -0.2667 | 0.2821 | 0.349698 |
| FUS BBBO+:30d | -1.1272 | 0.2821 | 0.000254 |
| Pairwise Differences of Treatment * Time with Tukey’s P value adjustment for a family of 6 estimates | | | |
| <u>Pair</u> | <u>Estimate</u> | <u>Std. Error</u> | <u>p-value</u> |
| (FUS BBBO- 1d) - (FUS BBBO+ 1d) | 0.1560 | 0.259 | 0.9902 |
| (FUS BBBO- 7d) - (FUS BBBO+ 7d) | 0.4228 | 0.259 | 0.5825 |
| (FUS BBBO- 30d) - (FUS BBBO+ 30d) | 1.2833 | 0.259 | 0.0002 |
| **Figure 6D: Linear Mixed Effect Model by Restricted Maximum Likelihood** | | | |

| lnVolume ~ Treatment * Time + Starting Volume + (1 \| Mouse) + (1 \| Lesion) | | | |
|---|---|---|---|
| Random Effects: | | | |
| Groups | Name | Variance | Std. Dev. |
| Lesion ID | (Intercept) | 0.42707 | 0.6535 |
| Mouse ID | (Intercept) | 0.07932 | 0.2816 |
| Residual | | 0.57282 | 0.7568 |
| Fixed Effects: | | | |
| Groups | Estimate | Std. Error | p-value |
| Intercept | -4.5358 | 0.2689 | 7.81E-16 |
| FUS BBBO+ | -0.4472 | 0.3061 | 0.147764 |
| 7d | 0.8683 | 0.2336 | 0.000362 |
| 30d | 1.658 | 0.2336 | 3.73E-10 |
| Starting Volume | 27.8133 | 4.1755 | 4.97E-08 |
| FUS BBBO+:7d | -0.7752 | 0.3231 | 0.018627 |
| FUS BBBO+:30d | -1.8455 | 0.3231 | 1.64E-07 |
| Pairwise Differences of Treatment * Time with Tukey's P value adjustment for a family of 6 estimates | | | |
| Pair | Estimate | Std. Error | p-value |
| (FUS BBBO- 1d) - (FUS BBBO+ 1d) | 0.4472 | 0.309 | 0.6973 |
| (FUS BBBO- 7d) - (FUS BBBO+ 7d) | 1.2224 | 0.309 | 0.0021 |
| (FUS BBBO- 30d) - (FUS BBBO+ 30d) | 2.2928 | 0.309 | <.0001 |
| **Figure 6F: Linear Mixed Effect Model by Restricted Maximum Likelihood** | | | |
| lnVolume ~ Treatment * Time + Starting Volume + (1 \| Mouse) + (1 \| Lesion) | | | |
| Random Effects: | | | |
| Groups | Name | Variance | Std. Dev. |
| Lesion ID | (Intercept) | 0.3568 | 0.5973 |
| Mouse ID | (Intercept) | 0 | 0 |
| Residual | | 0.3373 | 0.5808 |
| Fixed Effects: | | | |
| Groups | Estimate | Std. Error | p-value |
| Intercept | -3.860709 | 0.281417 | 4.12E-15 |
| FUS BBBO+ | -0.30401 | 0.364332 | 0.40962 |
| 7d | 0.178492 | 0.259738 | 0.49613 |
| 30d | 1.385354 | 0.259738 | 4.66E-06 |
| Starting Volume | 7.297587 | 1.269863 | 1.90E-05 |
| FUS BBBO+:7d | -0.006141 | 0.35888 | 0.98644 |
| FUS BBBO+:30d | -1.043723 | 0.35888 | 0.00604 |
| Pairwise Differences of Treatment * Time with Tukey's P value adjustment for a family of 6 estimates | | | |
| Pair | Estimate | Std. Error | p-value |
| (FUS BBBO- 1d) - (FUS BBBO+ 1d) | 0.3040 | 0.379 | 0.9651 |

| | | | |
|---|---|---|---|
| (FUS BBBO- 7d) - (FUS BBBO+ 7d) | 0.3102 | 0.379 | 0.9620 |
| (FUS BBBO- 30d) - (FUS BBBO+ 30d) | 1.3477 | 0.379 | 0.0131 |
| **Figure 7B: Wilcoxon Matched-Pairs Signed Rank Test** | | | |
| Comparison | | p-value | |
| Contra vs. BBBO | | 0.1250 | |
| **Figure 7D: Wilcoxon Matched-Pairs Signed Rank Test** | | | |
| Comparison | | p-value | |
| Contra vs. BBBO | | 0.0312 | |
| **Figure 7F: Wilcoxon Matched-Pairs Signed Rank Test** | | | |
| Comparison | | p-value | |
| Contra vs. BBBO | | 0.2500 | |
| **Figure 8B: Linear Mixed Effect Model by Restricted Maximum Likelihood** | | | |
| Avg *Krit1*KO Area ~ Treatment * Time + (1 \| Section) + (1 \| Mouse) | | | |
| Random Effects: | | | |
| Groups | Name | Variance | Std. Dev. |
| Section ID | (Intercept) | 13385 | 115.7 |
| Mouse ID | (Intercept) | 17059 | 130.6 |
| Residual | | 66274 | 257.4 |
| Fixed Effects: | | | |
| Groups | Estimate | Std. Error | p-value |
| Intercept | 226.086 | 114.556 | 0.07089 |
| FUS BBBO+ | 76.331 | 87.317 | 0.38320 |
| 7d | 225.378 | 149.651 | 0.16534 |
| 30d | 347.061 | 163.388 | 0.06444 |
| FUS BBBO+:7d | -65.911 | 107.298 | 0.53980 |
| FUS BBBO+:30d | -298.044 | 111.324 | 0.00813 |
| Pairwise Differences of Treatment * Time with Tukey's P value adjustment for a family of 6 estimates | | | |
| Pair | Estimate | Std. Error | p-value |
| (FUS BBBO- 1d) - (FUS BBBO+ 1d) | -76.3 | 88.2 | 0.9541 |
| (FUS BBBO- 7d) - (FUS BBBO+ 7d) | -10.4 | 63.1 | 1.0000 |
| (FUS BBBO- 30d) - (FUS BBBO+ 30d) | 221.7 | 69.2 | 0.0199 |
| **Figure 8D: Linear Mixed Effect Model by Restricted Maximum Likelihood** | | | |
| Iba1+ Count per $mm^2$ ~ Treatment * Time + (1 \| Section) + (1 \| Mouse) | | | |
| Random Effects: | | | |
| Groups | Name | Variance | Std. Dev. |
| Section ID | (Intercept) | 81311 | 285.2 |
| Mouse ID | (Intercept) | 1078220 | 1038.4 |
| Residual | | 480525 | 693.2 |
| Fixed Effects: | | | |

| Groups | Estimate | Std. Error | p-value |
|---|---|---|---|
| Intercept | 4467.453 | 627.892 | 0.000734 |
| FUS BBBO+ | -983.615 | 215.350 | 2.17e-05 |
| 7d | -1345.894 | 898.780 | 0.189967 |
| 30d | -3266.993 | 998.152 | 0.020358 |
| FUS BBBO+:7d | 1593.532 | 363.458 | 0.031262 |
| FUS BBBO+:30d | 808.586 | 367.456 | 0.00813 |
| Pairwise Differences of Treatment * Time with Tukey's P value adjustment for a family of 6 estimates | | | |
| Pair | Estimate | Std. Error | p-value |
| (FUS BBBO- 1d) - (FUS BBBO+ 1d) | 984 | 216 | 0.0003 |
| (FUS BBBO- 7d) - (FUS BBBO+ 7d) | -610 | 297 | 0.3209 |
| (FUS BBBO- 30d) - (FUS BBBO+ 30d) | 175 | 298 | 0.9916 |
| **Figure 8E: Linear Mixed Effect Model by Restricted Maximum Likelihood** | | | |
| ln(Avg Iba1+ Area) ~ Treatment * Time + (1 \| Section) + (1 \| Mouse) | | | |
| Random Effects: | | | |
| Groups | Name | Variance | Std. Dev. |
| Section ID | (Intercept) | 0.006646 | 0.08152 |
| Mouse ID | (Intercept) | 0.005774 | 0.07599 |
| Residual | | 0.087773 | 0.29627 |
| Fixed Effects: | | | |
| Groups | Estimate | Std. Error | p-value |
| Intercept | 3.42482 | 0.08566 | 1.39e-11 |
| FUS BBBO+ | 0.32206 | 0.09157 | 0.000791 |
| 7d | -0.40781 | 0.13213 | 0.009400 |
| 30d | -0.45037 | 0.14224 | 0.008849 |
| FUS BBBO+:7d | -0.02827 | 0.15271 | 0.853612 |
| FUS BBBO+:30d | -0.42245 | 0.15663 | 0.008899 |
| Pairwise Differences of Treatment * Time with Tukey's P value adjustment for a family of 6 estimates | | | |
| Pair | Estimate | Std. Error | p-value |
| (FUS BBBO- 1d) - (FUS BBBO+ 1d) | -0.3221 | 0.0922 | 0.0106 |
| (FUS BBBO- 7d) - (FUS BBBO+ 7d) | -0.2938 | 0.1250 | 0.1866 |
| (FUS BBBO- 30d) - (FUS BBBO+ 30d) | 0.1004 | 0.1273 | 0.9685 |
| **Figure 8G: Linear Mixed Effect Model by Restricted Maximum Likelihood** | | | |
| ln(CD68+ Cells per mm$^2$) ~ Treatment * Time + (1 \| Section) + (1 \| Mouse) | | | |
| Random Effects: | | | |
| Groups | Name | Variance | Std. Dev. |
| Section ID | (Intercept) | 0.9597 | 0.9796 |

| | | | |
|---|---|---|---|
| Mouse ID | (Intercept) | 3.5651 | 1.8882 |
| Residual | | 2.7935 | 1.6714 |
| Fixed Effects: | | | |
| Groups | Estimate | Std. Error | p-value |
| Intercept | 5.9312 | 1.2474 | 0.002629 |
| FUS BBBO+ | -2.3533 | 0.5727 | 6.15e-05 |
| 7d | 0.2004 | 1.7147 | 0.910966 |
| 30d | -0.4185 | 1.9019 | 0.833708 |
| FUS BBBO+:7d | 1.1625 | 0.7048 | 0.100854 |
| FUS BBBO+:30d | 2.7470 | 0.7279 | 0.000222 |
| Pairwise Differences of Treatment * Time with Tukey's P value adjustment for a family of 6 estimates | | | |
| Pair | Estimate | Std. Error | p-value |
| (FUS BBBO- 1d) - (FUS BBBO+ 1d) | 2.3533 | 0.576 | 0.0009 |
| (FUS BBBO- 7d) - (FUS BBBO+ 7d) | 1.1908 | 0.414 | 0.0507 |
| (FUS BBBO- 30d) - (FUS BBBO+ 30d) | -0.3937 | 0.450 | 0.9519 |
| **Figure 8H: Linear Mixed Effect Model by Restricted Maximum Likelihood** | | | |
| ln(% Ter119+ in Iba1+) ~ Treatment * Time + (1 \| Section) + (1 \| Mouse) | | | |
| Random Effects: | | | |
| Groups | Name | Variance | Std. Dev. |
| Section ID | (Intercept) | 0.07114 | 0.2667 |
| Mouse ID | (Intercept) | 0.47643 | 0.6902 |
| Residual | | 0.54384 | 0.7375 |
| Fixed Effects: | | | |
| Groups | Estimate | Std. Error | p-value |
| Intercept | 2.04628 | 0.46386 | 0.00265 |
| FUS BBBO+ | -0.07971 | 0.25005 | 0.75026 |
| 7d | 0.13130 | 0.63163 | 0.84174 |
| 30d | -0.96017 | 0.69874 | 0.21715 |
| FUS BBBO+:7d | -0.46773 | 0.30623 | 0.12842 |
| FUS BBBO+:30d | 0.15876 | 0.31861 | 0.61891 |
| Pairwise Differences of Treatment * Time with Tukey's P value adjustment for a family of 6 estimates | | | |
| Pair | Estimate | Std. Error | p-value |
| (FUS BBBO- 1d) - (FUS BBBO+ 1d) | 0.0797 | 0.252 | 0.9996 |
| (FUS BBBO- 7d) - (FUS BBBO+ 7d) | 0.5474 | 0.179 | 0.0303 |
| (FUS BBBO- 30d) - (FUS BBBO+ 30d) | -0.0790 | 0.198 | 0.9987 |
| **Figure S1B: Linear Mixed Effect Model by Restricted Maximum Likelihood** | | | |
| ln(% Ter119+ Area) ~ Treatment * Time + (1 \| Section) + (1 \| Mouse) | | | |

| Random Effects: | | | |
|---|---|---|---|
| Groups | Name | Variance | Std. Dev. |
| Section ID | (Intercept) | 0.34976 | 0.5914 |
| Mouse ID | (Intercept) | 0.07128 | 0.2670 |
| Residual | | 1.75762 | 1.3258 |
| Fixed Effects: | | | |
| Groups | Estimate | Std. Error | p-value |
| Intercept | 2.18802 | 0.46560 | 8.94e-05 |
| FUS BBBO+ | -0.08504 | 0.44544 | 0.8488 |
| 7d | -0.66803 | 0.57819 | 0.2657 |
| 30d | -1.40137 | 0.62072 | 0.0412 |
| FUS BBBO+:7d | -0.20092 | 0.54866 | 0.7146 |
| FUS BBBO+:30d | -0.08830 | 0.56995 | 0.8771 |
| Pairwise Differences of Treatment * Time with Tukey's P value adjustment for a family of 6 estimates | | | |
| Pair | Estimate | Std. Error | p-value |
| (FUS BBBO- 1d) - (FUS BBBO+ 1d) | 0.085 | 0.453 | 1.0000 |
| (FUS BBBO- 7d) - (FUS BBBO+ 7d) | 0.286 | 0.325 | 0.9507 |
| (FUS BBBO- 30d) - (FUS BBBO+ 30d) | 0.173 | 0.356 | 0.9966 |

| **Figure S2A: Linear Regression** | |
|---|---|
| Is slope significantly non-zero? | |
| F | 11.96 |
| DFn, DFd | 1, 30 |
| P value | 0.0017 |
| Equation | Y = -0.4156*X + 0.1854 |
| **Figure S2B: Linear Regression** | |
| Is slope significantly non-zero? | |
| F | 8.600 |
| DFn, DFd | 1, 30 |
| P value | 0.0064 |
| Equation | Y = -5.941*X + 3.982 |
| **Figure S2C: Linear Regression** | |
| Is slope significantly non-zero? | |
| F | 11.32 |
| DFn, DFd | 1, 30 |
| P value | 0.0021 |
| Equation | Y = -0.09265*X + 0.1796 |
| **Figure S2D: Linear Regression** | |
| Is slope significantly non-zero? | |
| F | 3.042 |
| DFn, DFd | 1, 30 |
| P value | 0.0914 |

| Equation | | Y = -0.8686*X + 3.471 | |
|---|---|---|---|
| **Figure S3A: Linear Mixed Effect Model by Restricted Maximum Likelihood** | | | |
| lnVol ~ Sex * Treatment * Time + StartVol + (1 \| Mouse) + (1 \| Lesion) + (1 \| Cohort) | | | |
| Random Effects: | | | |
| Groups | Name | Variance | Std. Dev. |
| Lesion ID | (Intercept) | 0.49951 | 0.7068 |
| Mouse ID | (Intercept) | 0.08626 | 0.2937 |
| Cohort | (Intercept) | 0.037 | 0.1924 |
| Residual | | 0.41209 | 0.6419 |
| Fixed Effects | | | |
| Groups | Estimate | Std. Error | p-value |
| (Intercept) | -4.43661 | 0.25911 | 3.44E-08 |
| M | 0.40544 | 0.36007 | 0.2659 |
| FUSBBBO+ | -0.02104 | 0.26514 | 0.9369 |
| 7d | 0.4201 | 0.18157 | 0.0219 |
| 30d | 1.11647 | 0.18157 | 5.50E-09 |
| Starting Volume | 10.56322 | 1.42424 | 1.20E-10 |
| M:FUSBBBO+ | -0.52574 | 0.42373 | 0.2168 |
| M:7d | 0.33961 | 0.29065 | 0.2443 |
| M:30d | 0.80099 | 0.29065 | 0.0065 |
| FUS BBBO+:7d | -0.40329 | 0.2498 | 0.1083 |
| FUS BBBO+:30d | -1.28541 | 0.2498 | 7.37E-07 |
| M:FUSBBBO+:7d | -0.16299 | 0.3966 | 0.6816 |
| M:FUSBBBO+:30d | -0.46596 | 0.3966 | 0.2417 |
| Pairwise Differences of Treatment * Time * Sex with Tukey's P value adjustment for a family of 12 estimates | | | |
| Pair | Estimate | Std. Error | p-value |
| (FUS BBBO- 1d F) - (FUS BBBO+ 1d F) | 0.02104 | 0.267 | 1.0000 |
| (FUS BBBO- 1d F) - (FUS BBBO- 1d M) | -0.40544 | 0.394 | 0.9961 |
| (FUS BBBO+ 1d F) - (FUS BBBO+ 1d M) | 0.12029 | 0.357 | 1.0000 |
| (FUS BBBO- 1d M) - (FUS BBBO+ 1d M) | 0.54677 | 0.332 | 0.8881 |
| (FUS BBBO- 7d F) - (FUS BBBO+ 7d F) | 0.42433 | 0.267 | 0.9094 |
| (FUS BBBO- 7d F) - (FUS BBBO- 7d M) | -0.74505 | 0.394 | 0.7596 |
| (FUS BBBO+ 7d F) - (FUS BBBO+ 7d M) | -0.05632 | 0.357 | 1.0000 |
| (FUS BBBO- 7d M) - (FUS BBBO+ 7d M) | 1.11305 | 0.332 | 0.0460 |

| | | | |
|---|---|---|---|
| (FUS BBBO- 30d F) - (FUS BBBO+ 30d F) | 1.30645 | 0.267 | 0.0002 |
| (FUS BBBO- 30d F) - (FUS BBBO- 30d M) | -1.20643 | 0.394 | 0.1253 |
| (FUS BBBO+ 30d F) - (FUS BBBO+ 30d M) | -0.21473 | 0.357 | 1.0000 |
| (FUS BBBO- 30d M) - (FUS BBBO+ 30d M) | 2.29814 | 0.332 | <.0001 |
| **Figure S3C: Linear Mixed Effect Model by Restricted Maximum Likelihood** | | | |
| De Novo Lesions ~ Sex * Treatment + StartingVolume + (1 \| Mouse) + (1 \| Cohort) | | | |
| Random Effects: | | | |
| Groups | Name | Variance | Std. Dev. |
| Mouse ID | (Intercept) | 1.664 | 1.29 |
| Cohort | (Intercept) | 9.58 | 3.095 |
| Residual | | 1.715 | 1.31 |
| Fixed Effects: | | | |
| Groups | Estimate | Std. Error | p-value |
| (Intercept) | 4.6864 | 1.9186 | 0.12436 |
| M | -0.5358 | 1.0766 | 0.62392 |
| FUS BBBO+ | -2.1063 | 0.608 | 0.0036 |
| Starting Volume | -9.7962 | 3.4981 | 0.00972 |
| M:FUS BBBO+ | 1.6714 | 0.9845 | 0.11182 |
| Pairwise Differences of Treatment * Sex with Tukey's P value adjustment for a family of 4 estimates | | | |
| Pair | Estimate | Std. Error | p-value |
| (FUS BBBO- F) - (FUS BBBO+ F) | 2.106 | 0.615 | 0.0198 |
| (FUS BBBO- M) - (FUS BBBO+ M) | 0.435 | 0.809 | 0.9483 |
| (FUS BBBO- F) - (FUS BBBO- M) | 0.536 | 1.107 | 0.9618 |
| (FUS BBBO+ F) - (FUS BBBO+ M) | -1.136 | 1.044 | 0.7009 |
| **Figure S4B: Linear Mixed Effect Model by Restricted Maximum Likelihood** | | | |
| ln(Ki67+*Krit1*KO+ per $mm^2$) ~ Treatment * Time + (1 \| Section) + (1 \| Mouse) | | | |
| Random Effects: | | | |
| Groups | Name | Variance | Std. Dev. |
| Section ID | (Intercept) | 0.2263 | 0.4757 |
| Mouse ID | (Intercept) | 2.6543 | 1.6292 |
| Residual | | 0.9272 | 0.9629 |
| Fixed Effects: | | | |
| Groups | Estimate | Std. Error | p-value |
| Intercept | 2.880500 | 1.064877 | 0.0202 |
| FUS BBBO+ | -0.009568 | 0.581157 | 0.9869 |
| 7d | 0.182283 | 1.472652 | 0.9039 |

| | | | |
|---|---|---|---|
| 30d | -1.173005 | 1.310476 | 0.3916 |
| FUS BBBO+:7d | 0.008108 | 0.737059 | 0.9913 |
| FUS BBBO+:30d | -0.100237 | 0.636427 | 0.8752 |
| Pairwise Differences of Treatment * Time with Tukey's P value adjustment for a family of 6 estimates | | | |
| Pair | Estimate | Std. Error | p-value |
| (FUS BBBO- 1d) - (FUS BBBO+ 1d) | 0.00957 | 0.589 | 1.0000 |
| (FUS BBBO- 7d) - (FUS BBBO+ 7d) | 0.00146 | 0.456 | 1.0000 |
| (FUS BBBO- 30d) - (FUS BBBO+ 30d) | 0.10981 | 0.261 | 0.9982 |
| **Figure S4C: Linear Mixed Effect Model by Restricted Maximum Likelihood** | | | |
| ln(Ki67+Iba1+ per mm$^2$) ~ Treatment * Time + (1 \| Section) + (1 \| Mouse) | | | |
| Random Effects: | | | |
| Groups | Name | Variance | Std. Dev. |
| Section ID | (Intercept) | 0.1474 | 0.384 |
| Mouse ID | (Intercept) | 2.5355 | 1.592 |
| Residual | | 1.5007 | 1.225 |
| Fixed Effects: | | | |
| Groups | Estimate | Std. Error | p-value |
| Intercept | 3.5181 | 1.0977 | 0.00686 |
| FUS BBBO+ | -0.5853 | 0.7126 | 0.41391 |
| 7d | -0.0327 | 1.5031 | 0.98302 |
| 30d | -1.8682 | 1.3348 | 0.18863 |
| FUS BBBO+:7d | 0.6657 | 0.9125 | 0.46786 |
| FUS BBBO+:30d | 0.5988 | 0.7846 | 0.44758 |
| Pairwise Differences of Treatment * Time with Tukey's P value adjustment for a family of 6 estimates | | | |
| Pair | Estimate | Std. Error | p-value |
| (FUS BBBO- 1d) - (FUS BBBO+ 1d) | 0.5853 | 0.721 | 0.9645 |
| (FUS BBBO- 7d) - (FUS BBBO+ 7d) | -0.0804 | 0.573 | 1.0000 |
| (FUS BBBO- 30d) - (FUS BBBO+ 30d) | -0.0135 | 0.330 | 1.0000 |
| **Figure S4D: Linear Mixed Effect Model by Restricted Maximum Likelihood** | | | |
| ln(Ki67+GFAP+ per mm$^2$) ~ Treatment * Time + (1 \| Section) + (1 \| Mouse) | | | |
| Random Effects: | | | |
| Groups | Name | Variance | Std. Dev. |
| Section ID | (Intercept) | 0.3824 | 0.6184 |
| Mouse ID | (Intercept) | 1.7387 | 1.3186 |
| Residual | | 2.0025 | 1.4151 |
| Fixed Effects: | | | |
| Groups | Estimate | Std. Error | p-value |

| | | | |
|---|---|---|---|
| Intercept | 3.48804 | 1.04563 | 0.00368 |
| FUS BBBO+ | -0.20148 | 0.83931 | 0.81089 |
| 7d | 0.62167 | 1.40384 | 0.66436 |
| 30d | -1.72514 | 1.24363 | 0.18655 |
| FUS BBBO+:7d | -0.62324 | 1.06992 | 0.56186 |
| FUS BBBO+:30d | -0.05176 | 0.92144 | 0.95534 |
| Pairwise Differences of Treatment * Time with Tukey's P value adjustment for a family of 6 estimates | | | |
| Pair | Estimate | Std. Error | p-value |
| (FUS BBBO- 1d) - (FUS BBBO+ 1d) | 0.20148 | 0.852 | 0.9999 |
| (FUS BBBO- 7d) - (FUS BBBO+ 7d) | 0.82472 | 0.667 | 0.8177 |
| (FUS BBBO- 30d) - (FUS BBBO+ 30d) | 0.25324 | 0.383 | 0.9854 |
| **Figure S4E: Linear Mixed Effect Model by Restricted Maximum Likelihood** | | | |
| ln(GFAP+ Cells per $mm^2$) ~ Treatment * Time + (1 \| Section) + (1 \| Mouse) | | | |
| Random Effects: | | | |
| Groups | Name | Variance | Std. Dev. |
| Section ID | (Intercept) | 880.3 | 29.67 |
| Mouse ID | (Intercept) | 1777.2 | 42.16 |
| Residual | | 36136.3 | 190.10 |
| Fixed Effects: | | | |
| Groups | Estimate | Std. Error | p-value |
| Intercept | 554.270 | 90.530 | 8.22e-08 |
| FUS BBBO+ | -64.152 | 104.608 | 0.541 |
| 7d | 7.178 | 112.619 | 0.949 |
| 30d | -165.920 | 99.670 | 0.103 |
| FUS BBBO+:7d | 1.594 | 134.986 | 0.991 |
| FUS BBBO+:30d | 53.108 | 116.075 | 0.648 |
| Pairwise Differences of Treatment * Time with Tukey's P value adjustment for a family of 6 estimates | | | |
| Pair | Estimate | Std. Error | p-value |
| (FUS BBBO- 1d) - (FUS BBBO+ 1d) | 64.15 | 106.4 | 0.9906 |
| (FUS BBBO- 7d) - (FUS BBBO+ 7d) | 62.56 | 85.7 | 0.9776 |
| (FUS BBBO- 30d) - (FUS BBBO+ 30d) | 11.04 | 51.0 | 0.9999 |
| **Figure S4F: Linear Mixed Effect Model by Restricted Maximum Likelihood** | | | |
| ln(Avg GFAP+ Area) ~ Treatment * Time + (1 \| Section) + (1 \| Mouse) | | | |
| Random Effects: | | | |
| Groups | Name | Variance | Std. Dev. |
| Section ID | (Intercept) | 0.03284 | 0.1812 |
| Mouse ID | (Intercept) | 0.07592 | 0.2755 |

| | | | |
|---|---|---|---|
| Residual | | 0.19686 | 0.4437 |
| Fixed Effects: | | | |
| <u>Groups</u> | <u>Estimate</u> | <u>Std. Error</u> | <u>p-value</u> |
| Intercept | 2.672961 | 0.272736 | 4.62e-10 |
| FUS BBBO+ | -0.100352 | 0.259908 | 0.700 |
| 7d | 0.216861 | 0.356642 | 0.551 |
| 30d | 0.030386 | 0.315486 | 0.924 |
| FUS BBBO+:7d | 0.007726 | 0.329501 | 0.981 |
| FUS BBBO+:30d | 0.122625 | 0.285849 | 0.669 |
| Pairwise Differences of Treatment * Time with Tukey’s P value adjustment for a family of 6 estimates | | | |
| <u>Pair</u> | <u>Estimate</u> | <u>Std. Error</u> | <u>p-value</u> |
| (FUS BBBO- 1d) - (FUS BBBO+ 1d) | 0.1004 | 0.264 | 0.9989 |
| (FUS BBBO- 7d) - (FUS BBBO+ 7d) | 0.0926 | 0.204 | 0.9975 |
| (FUS BBBO- 30d) - (FUS BBBO+ 30d) | -0.0223 | 0.120 | 1.0000 |
| **Figure S5B: Linear Mixed Effect Model by Restricted Maximum Likelihood** | | | |
| ln(CD45+ Cells per mm$^2$) ~ Treatment * Time + (1 \| Section) + (1 \| Mouse) | | | |
| Random Effects: | | | |
| <u>Groups</u> | <u>Name</u> | <u>Variance</u> | <u>Std. Dev.</u> |
| Section ID | (Intercept) | 0.1683 | 0.4103 |
| Mouse ID | (Intercept) | 0.4372 | 0.6612 |
| Residual | | 1.2472 | 1.1168 |
| Fixed Effects: | | | |
| <u>Groups</u> | <u>Estimate</u> | <u>Std. Error</u> | <u>p-value</u> |
| Intercept | 4.4152 | 0.4809 | 6.5e-05 |
| FUS BBBO+ | 0.5764 | 0.3464 | 0.1011 |
| 7d | -0.7417 | 0.7132 | 0.3303 |
| 30d | -1.5646 | 0.7782 | 0.0842 |
| FUS BBBO+:7d | 0.8158 | 0.5824 | 0.1656 |
| FUS BBBO+:30d | -1.5080 | 0.5915 | 0.0133 |
| Pairwise Differences of Treatment * Time with Tukey’s P value adjustment for a family of 6 estimates | | | |
| <u>Pair</u> | <u>Estimate</u> | <u>Std. Error</u> | <u>p-value</u> |
| (FUS BBBO- 1d) - (FUS BBBO+ 1d) | -0.576 | 0.348 | 0.5647 |
| (FUS BBBO- 7d) - (FUS BBBO+ 7d) | -1.392 | 0.476 | 0.0494 |
| (FUS BBBO- 30d) - (FUS BBBO+ 30d) | 0.932 | 0.480 | 0.3882 |
